# Heterogeneous but not random: Cargo degradation in phagosomes is kinetically coupled to their intracellular mobility

**DOI:** 10.1101/2021.04.04.438376

**Authors:** Yanqi Yu, Zihan Zhang, Glenn F. W. Walpole, Yan Yu

## Abstract

Immune cells degrade internalized pathogens in phagosomes through sequential biochemical changes. The degradation must be fast enough for effective infection control. The presumption is that each phagosome degrades cargos autonomously with a distinct but stochastic kinetic rate. Here we report that the degradation kinetics of individual phagosomes is not stochastic but coupled to their intracellular motility. By engineering RotSensors that are optically anisotropic, magnetic responsive, and fluorogenic in response to degradation activities in phagosomes, we monitored cargo degradation kinetics in single phagosomes simultaneously with their translational and rotational dynamics. We show that phagosomes that move faster centripetally are more likely to encounter and fuse with lysosomes, thereby acidifying faster and degrading cargos more efficiently. The degradation rates increase nearly linearly with the translational and rotational velocities of phagosomes. Our results indicate that the centripetal motion of phagosomes functions as a clock for controlling the progression of cargo degradation.

## Introduction

Innate immune cells, such as macrophages, ingest pathogens and dead cells via a phagocytosis process. This is an essential process in infection control and tissue homeostasis. After recognizing ligands on pathogens, immune cells internalize them into phagosomes and transport them from the cell periphery to the perinuclear region^1, 2, 3, 4, 5^. During this intracellular transport, phagosomes degrade the entrapped pathogens through sequential biochemical transformations, including the acidification of the lumen^6^, the activation of hydrolytic enzymes^7^, and the generation of reactive oxygen species (ROS)^8^. Phagosomes must degrade pathogens efficiently to prevent them from escaping. Many pathogens, such as the bacterium *Legionella pneumophila*, were found to evade the degradative environment in phagosomes by hijacking the intracellular trafficking process^9, 10, 11^. These examples highlight that the degradative function of phagosomes is tightly associated with their dynamics in cells. Understanding the relationship between the degradative function and dynamics of phagosomes is critical for developing therapeutics against infections and inflammation. It is, however, challenging due to the autonomous nature of the two processes. Each phagosome degrades its cargo at its own distinct kinetic rate, which was assumed to be stochastic. The velocity at which phagosomes are transported along microtubules is also heterogeneous and believed to be stochastic^2, 12^. Therefore, those dynamic and biochemical changes must be quantified at the single phagosome level to reveal their relationships, but there is currently no method to do so. The dynamics of phagosomes in cells has been extensively studied using single-particle imaging^13, 14^. The degradative activities, including lumen acidification and protein digestion, in phagosomes have been studied separately in population-based measurements^15^. In those ensemble-average studies, the individuality of phagosomes was overlooked. Regarding the relationship between the degradative function and dynamics of phagosomes, studies have relied on approaches that inhibit phagosome dynamics by disrupting the cytoskeleton or molecular motor functions. The disruption of actin was shown to attenuate phagosome-endosome fusion^16, 17, 18^. Microtubule depolymerization was found to disrupt: i) phagosome-lysosome fusion^19^, ii) content delivery from late endosomes to phagosomes^20^, and iii) acquisition of lysosome markers^21^. These observations demonstrated that dynamics of phagosomes in cells is important for their degradative function, but none of those studies identified the quantitative relationship between phagosome dynamics and kinetics of the biochemical activities during cargo degradation.

In this study, we develop a single-phagosome imaging toolset for monitoring both degradative function and dynamics of individual phagosomes. The RotSensor particles are optically anisotropic and fluorogenic in response to acidification in phagosomes and phagosome-lysosome fusion, two key degradative activities. Using the RotSensors as cargos in phagosomes, we imaged simultaneously the biochemical changes during phagosome degradation and their translational and rotational dynamics in living cells. By combining this toolset with magnetic tweezers to control the dynamics of phagosomes, we obtained direct evidence that the microtubule-based centripetal transport of phagosomes determines the kinetic rates of their degradation process. More mobile phagosomes fuse with lysosomes and acidify in their lumen more rapidly, following a nearly linear relationship. Our results reveal the orchestration of the dynamical and biochemical processes that occur during pathogen degradation in phagosomes, providing a possible mechanism by which some pathogens may evade immune clearance by disrupting the intracellular dynamics of residing phagosomes.

## Results

### pH-responsive rotational particle sensors (pH-RotSensors) for measuring phagosome acidification and dynamics

The acidification of phagosome lumen is an early indicator of its degradative function and required for many subsequent events, including the activation of degradative enzymes^22, 23^. Therefore, we first designed pH-responsive RotSensors to simultaneously measure the acidification of single phagosomes and their intracellular transport dynamics, including both translational and rotational movements (Fig. 1a). Each RotSensor consists of a pH-responsive particle (1 μm diameter) covalently conjugated with a smaller green particle (100 nm diameter) with non-overlapping fluorescence emission (Fig. 1b). Once engulfed into phagosomes, the “snowman”-like RotSensors report both the translational and the rotational motion of the residing phagosome. The azimuthal (φ) and polar (θ) angles of each RotSensor can be measured, respectively, from the orientation and projection distance between the pair of the two particles, as previously described (Fig. 1b)^24^.

**Fig. 1.**
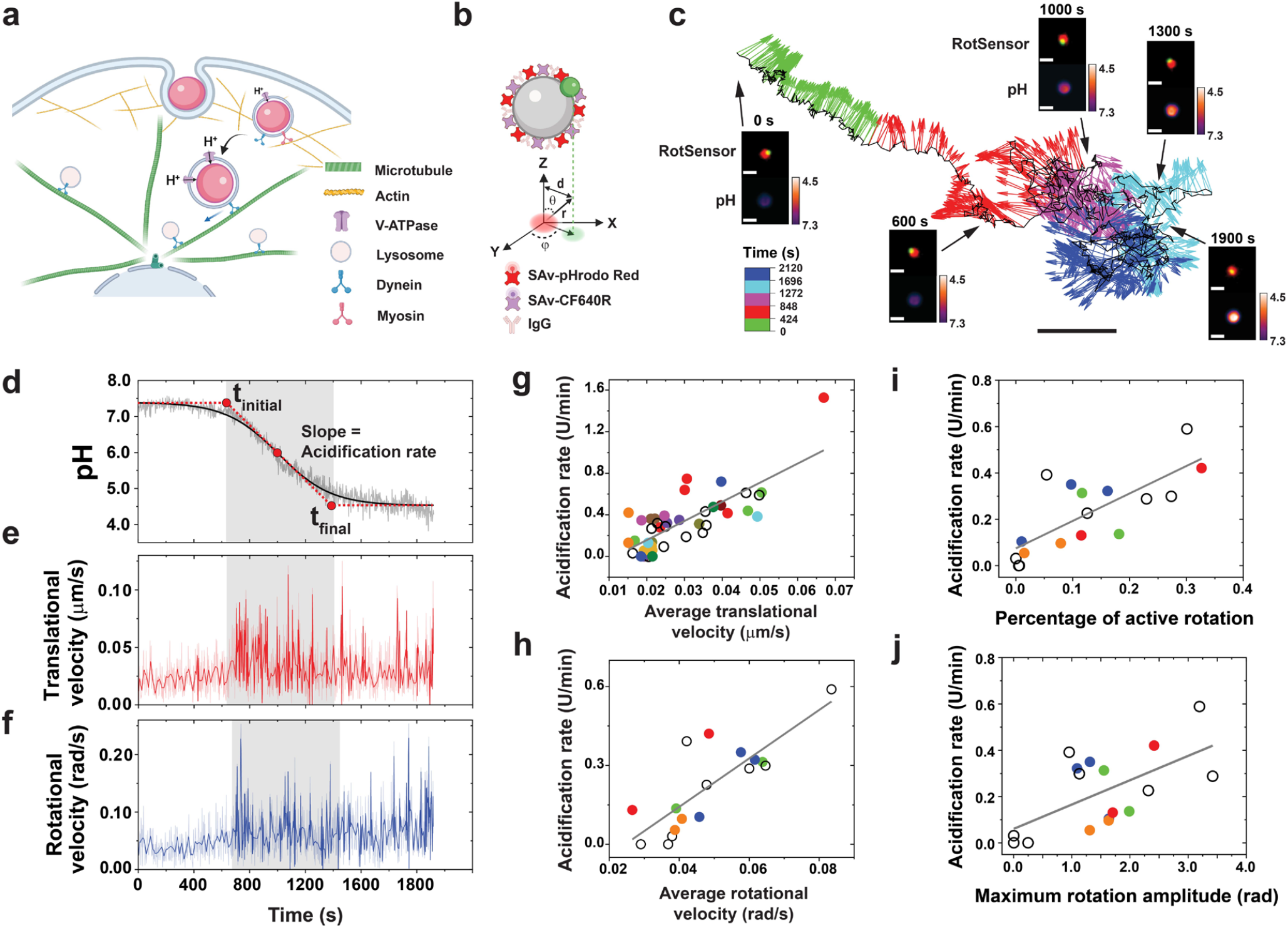
Simultaneous measurement of phagosome dynamics and acidification. **a** Schematics showing internalization of pH-sensitive RotSensors into phagosomes in macrophage cells. **b** Schematic illustration of the RotSensor design. Each RotSensor contains a 1 µm silica particle tethered with a 100 nm yellow-green fluorescent particle. The silica particles were coated with streptavidin (SAv)-pHrodo Red (pH reporter) and SAv-CF640R (reference). The azimuthal (φ) and polar (θ) angles of each pH-RotSensor were analyzed based on its projection fluorescence image. d and r denote the projection inter-particle distance and the physical inter-particle distance, respectively. **c** A representative trajectory of a pH-RotSensor-containing phagosome. Vectors indicates the orientation of the pH-RotSensor and are color-coded based on time. Scale bar, 2 µm. Insets show fluorescence images of the RotSensor (top) and the pH response (bottom). Scale bar, 1 µm. **d-f** Line plots showing acidification (**d**), translational (**e**) and rotational velocities (**f**) of the pH-RotSensor-containing phagosome in **c**. The acidification profile is fitted with sigmoidal-Boltzmann function to determine the initial pH, final pH, the period of rapid acidification (t_initial_ to t_final_), and acidification rate. The period of rapid acidification is highlighted in grey. Darker lines in **e** and **f** are velocity values after wavelet denoising. **g** and **h** Scattered plots showing acidification rates against translational (**g**) and rotational velocities (**h**) of single phagosomes. Each data point represents one single phagosome. Data points from multiple phagosomes within the same cells are shown in the same solid color. Data points from cells containing only one phagosome are shown as black circles. N = 42 phagosomes from 24 cells for translational tracking, and 17 phagosomes from 12 cells for rotational tracking. Pearson’s coefficients of 0.78 and 0.81 were obtained in **g** and **h**, respectively. **I** and **j** Scatter plots showing acidification rates against percentage of active rotation (**i**) and maximum rotation amplitude (**j**) of single phagosomes during rapid acidification period. N = 17 phagosomes from 12 cells. Pearson’s coefficients of 0.76 and 0.61 are obtained in **i** and **j**.

To monitor phagosome acidification, the 1 μm particle in each RotSensor was biotinylated and subsequently conjugated with two types of streptavidin, one type labeled with the pH-indicator pHrodo Red and the other with the reference dye CF640R. The dye pHrodo Red fluoresces intensely in an acidic environment, making it an ideal indicator of phagosome acidification. CF640R was chosen as a reference dye because it is pH insensitive and photostable (Supplementary Fig. 1). Streptavidin is critical in the RotSensor design because it acts as a cushion layer to separate the pHrodo Red dye from the particle surface. Without the streptavidin linker, pHrodo Red that was directly conjugated onto the particle surface exhibited little pH sensitivity. The pH-sensitive RotSensors were then coated with immunoglobulin G (IgG) via non-specific adsorption to initiate Fc receptor (FcR)-dependent phagocytosis in RAW264.7 macrophage cells.

To quantify the pH response of the RotSensors, we measured the ratio of fluorescence emission intensities of pHrodo Red and of the reference dye CF640R; *I*_*pHrodo*_*/I*_*ref*_, as we varied pH inside phagosomes in live cells and in aqueous buffers (Supplementary Fig. 2). For both cases, *I*_*pHrodo*_*/I*_*ref*_ of the RotSensors increased linearly with decreasing pH, consistent with previous reports^25^. The pH responses of individual RotSensors, including the initial *I*_*pHrodo*_*/I*_*ref*_ ratio and slope, varied slightly from one to another, because of the different amounts of dyes on particles and the slightly different phagosome lumen environments^26, 27, 28, 29^. To eliminate the effect of such variation in pH measurements, we calibrated our pH measurements for each phagosome. We measured *I*_*pHrodo*_/*I*_*ref*_ of each phagosome versus time and then converted the intensity ratio to pH based on *I*_*pHrodo*_ /*I*_*ref*_ of the same phagosome at pH 4.5 and 7.3 (details in Methods section).

### pH-RotSensors reveal correlation between phagosome acidification rate and transport dynamics

Before tracking the dynamics of phagosomes, we first asked whether the rotation of RotSensors faithfully reports the rotation of their residing phagosomes. Transmission electron microscopy (TEM) images show that ≈ 86% of internalized RotSensors remained tightly wrapped by the phagosome membrane at the end of our experiments (50 min after particles were added to cells) (Supplementary Fig. 3). Based on the tight membrane fitting and the presence of ligand-receptor binding between the RotSensors and phagosome membranes, the RotSensors are unlikely to freely rotate inside phagosomes. Thus, their rotation is expected to report the rotation of their residing phagosomes.

We then simultaneously imaged the pH response and the rotational and translational dynamics of single phagosomes encapsulating RotSensors (Fig. 1c). Phagosomes acidified in three steps: 1) an initial standby period during which the phagosome pH remained at neutral pH of ≈ 7.3 (Fig. 1d), 2) a rapid acidification period when the pH quickly dropped over a period of a few minutes, and 3) an eventually plateau at pH 4.5-5.0. This acidic pH is a prerequisite for activating degradative enzymes^8, 30^. While individual phagosomes reached slightly different final pH values, their acidification kinetics mostly (31 of 42 phagosomes) followed a sigmoidal-Boltzmann function:

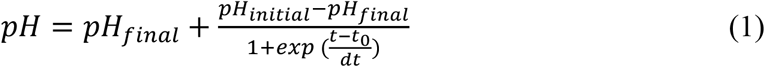

in which *t*_0_ is the half-response point, and *t*_*initial*_ and *t*_*final*_ denote the beginning and the end of rapid acidification, respectively (Fig. 1d). Slope at *t*_0_ is the acidification rate by definition. We found that phagosomes acidified at an average rate of 0.33 ± 0.28 pH unit/min (N = 42). The acidification of phagosomes requires intact microtubules, as the acidification was abolished upon microtubule disruption by nocodazole (Supplementary Fig. 4).

For transport dynamics, we observed that a majority (∼70%) of phagosomes exhibited a slow-to-fast movement transition both translationally and rotationally (Fig. 1e and f). The remaining 30% of phagosomes exhibited only slow movements. This initially slow movements are not dependent on microtubules, as the average velocities were not affected by microtubule depolymerization (Supplementary Fig. 5). After a few minutes of slow movements, phagosomes moved rapidly centripetally with velocities as high as 0.1 µm/s (Fig. 1e). This range of phagosome velocities is consistent with literature reports for 1 µm bead-containing phagosomes in macrophage cells^13^. Meanwhile, phagosomes also rotated frequently in between segments of translational motion with rotational velocities up to 0.15 rad/s (Fig. 1f). These dynamic behaviors, particularly phagosome rotation, are characteristic of cargo transport driven by microtubule-based molecular motors, as we previously reported^31^. This was further confirmed in control experiments, in which the rapid translational and rotational dynamics disappeared in cells treated with nocodazole (Supplementary Fig. 4).

Notably, the beginning of the rapid acidification period appeared to coincide with the onset of fast microtubule-based transport (Fig. 1e and f). To explore the possible correlation between the two processes, we calculated the acidification rate of single phagosomes based on the sigmoidal fitting of their individual pH-time plots (Fig. 1d). We then plotted the acidification rate of single phagosomes against their individual translational and rotational velocities during the rapid acidification phase (from *t*_*initial*_ to *t*_*final*_) (Fig. 1g and h). The acidification rate varied broadly among phagosomes, even those from the same cells. This reflected the individuality of each phagosome in degradation. Despite this heterogeneity, the acidification rates of individual phagosomes follow a linear relationship with both their translational and rotational velocities with a Pearson’s coefficient of 0.78 and 0.81, respectively. Those observations apply not only to phagosomes from different cells but also to those from the same cells. This is shown in Fig. 1g and h, in which data of phagosomes from the same cells are the same color. This is also a general phenomenon independent of the activation status of macrophage cells (Supplementary Fig. 6).

Further, the acidification rates of single phagosomes also follow a linear relationship with the percentage, maximum amplitude, and frequency of active rotation of phagosomes during the rapid acidification phase (from *t*_*initial*_ to *t*_*final*_) with a Pearson’s coefficient of 0.76, 0.60 and 0.61, respectively (Fig. 1i and j, and Supplementary Fig. 7). For this analysis, active rotation of phagosomes was distinguished from passive random rotation by using a Wavelet analysis method reported previously (Supplementary Fig. 8; Methods)^32^. The percentage of active rotation was defined as the total time during which the phagosome underwent active rotation divided by the time duration of its rapid acidification (from *t*_*initial*_ to *t*_*final*_). The results altogether indicate that more mobile phagosomes acidify faster. As an additional evidence supporting this conclusion, we also observed that the ∼30% of phagosomes that never underwent rapid microtubule-based transport exhibited minimal acidification (Supplementary Fig. 19a-c). Surprisingly however, the final pH attained by individual phagosomes showed no correlation with either the translational or the rotational velocities of those phagosomes (Supplementary Fig. 10). Because the final acidic pH determines whether a phagosome can successfully degrade its cargo, this result suggests that the degradative capacity of a phagosome is likely determined prior to its transport on microtubules, but that the rate of degradation is correlated with the transport dynamics.

### FRET-RotSensors for measuring phagosome-lysosome fusion

During cargo degradation, phagosome acidification relies on its continuous fusion with lysosomes, which deliver proton-pumping vacuolar H+-ATPase (V-ATPase) to it^28, 33, 34, 35, 36^. Since we found that more mobile phagosomes acidify more rapidly, we asked whether phagosome-lysosome fusion is likewise correlated with phagosome mobility. To do this, we measured the kinetics of phagosome-lysosome fusion by modifying the RotSensor design into a Förster Resonance Energy Transfer (FRET)-based assay. This fluorescence assay can be used to determine when two fluorophores, a donor inside lysosomes and an acceptor inside phagosomes, are in close proximity to one another. FRET-sensitive RotSensors were prepared in the same way as pH-sensitive RotSensors, but, in this case, the sensor particles were labeled with streptavidin that was conjugated with the donor fluorophore Alexa568 (Fig. 2a). For FRET, lysosomes were loaded with fluid-phase biotinylated bovine serum albumin (BSA) that was conjugated with the acceptor fluorophore Alexa647 using a pulse–chase procedure^37^. We confirmed that the BSA-Alexa647-biotin was contained inside lysosomes based on its colocalization with the lysosome marker LysoTracker inside cells (Supplementary Fig. 11). Inside lysosomes, BSA-Alexa647-biotin was fragmented by proteases before RotSensors were added to cells for internalization (Supplementary Fig. 12). The working principle of the FRET assay is that once the contents of lysosomes are intermixed with the phagosome lumen during fusion, the fragmented BSA-Alexa647-biotin from lysosomes will in close proximity with Alexa568 conjugated RotSensors leading to FRET. The magnitude of the FRET signal thus indicates the extent of phagosome-lysosome fusion. This assay doesn’t necessarily require biotin-streptavidin binding. We showed in *in vitro* experiments that FRET can occur when BSA-Alexa647, without biotinylation, was added to FRET-RotSensors in 1xPBS solution (Supplementary Fig. 13). However, we still used BSA-Alexa647-biotin in this FRET assay because it was loaded into lysosomes much more efficiently than BSA-Alexa647 alone, which improved the sensitivity of the FRET assay (Supplementary Fig. 14). The observation that the biotin moiety promotes uptake of BSA by cells is consistent with previous reports^38, 39^.

**Fig. 2.**
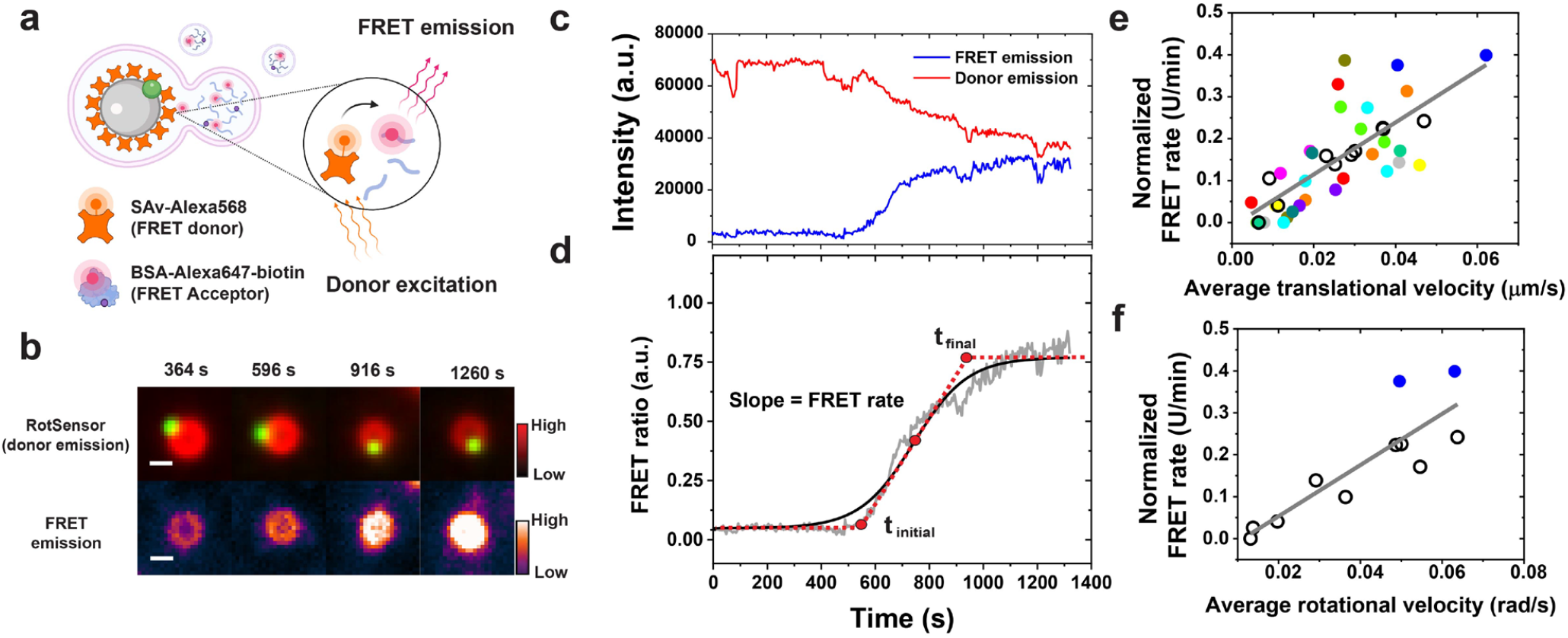
Simultaneous measurement of phagosome-lysosome fusion and phagosome transport dynamics. **a** Schematic illustration of the Förster resonance energy transfer (FRET)-based phagosome-lysosome fusion assay. The FRET-RotSensor was coated with SAv-Alexa568 (FRET donor). BSA-Alexa647-biotin (FRET acceptor) was loaded into lysosomes. Phagosome fusion with lysosomes leads to intermixing between donor fluorophore (Alexa568) and fluid phase acceptor fluorophore (Alexa647) generating FRET emission (680 nm) under the donor excitation of 561 nm. **b** and **c** Fluorescence images and intensity plots showing the change of FRET emission (ex/em: 561/680 nm) and donor emission (red, ex/em: 561/586 nm) from a FRET-RotSensor (100 nm yellow-green nanoparticle in green) in a RAW264.7 macrophage. Scale bar, 2μm. **d** FRET ratio vs. time plot is fitted with sigmoidal-Boltzmann function (shown as the black solid line) to determine the initial time point of phagosome-lysosome fusion (t_initial_), the time point where FRET-signal reaches a plateau (t_final_), and the FRET rate, as indicated by the red dotted line. **e** and **f** Scatter plots showing normalized FRET rate against translational (**e**) and rotational velocities (**f**) of single phagosomes during the period of its fusion with lysosomes. Each data point represents data from a single phagosome. Data points from multiple phagosomes within the same cells are shown in the same solid color. Data points from cells containing only one phagosome are shown as black circles. For translational tracking, N = 40 phagosomes from 22 cells. For rotational tracking, N = 11 phagosomes from 10 cells. The black lines indicate linear regression with a Pearson’s coefficient of 0.75 in **e** and of 0.86 in **f**.

For a thorough validation of the FRET assay, we first tested the FRET-RotSensors *in vitro* by mixing them with BSA-Alexa647 in 1xPBS (Supplementary Fig. 13). In this experiment, we could not measure the acceptor emission in the FRET channel due to the excessive amount of free BSA-Alexa647 in solution. Instead, we quantified the percentage of donor quenching as an indirect indicator of FRET efficiency. We found that the percentage of donor quenching, first increased linearly as a function of BSA-Alexa647 concentration in the lysosome loading solution ([BSA-Alexa647]_loading_), but reached a plateau at [BSA-Alexa647]_loading_ > 200 µg/ml. This plateau indicates the maximum FRET efficiency which is determined by the amount of the donor fluorophore (Alexa568) available on RotSensors. For the FRET assay to accurately detect continuous fusion of a phagosome with lysosomes, the concentration of acceptor fluorophores (Alexa647) loaded into lysosomes must be optimized to achieve sufficient FRET efficiency without saturating the donor on the RotSensors. We did this by optimizing the loading concentration of BSA-Alexa647-biotin into lysosomes.

We next tested and optimized the FRET assay in live cells. To start with, we loaded lysosomes using 10 µg/mL BSA-Alexa647-biotin ([BSA-Alexa647-biotin]_loading_ = 10 µg/mL). After internalization of FRET-RotSensors, lysosomes docked on the phagosomes, and the FRET-RotSensors inside phagosomes exhibited decreased fluorescence emission in the donor channel (ex/em: 561/586 nm) and simultaneously increased emission in FRET channel (ex/em: 561/680 nm) (Fig. 2b and c). For quantification, we calculated FRET ratio defined as:

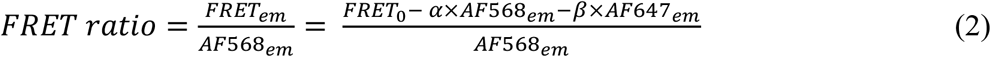

In this equation, *FTET*_0_ is the measured FRET signal (ex/em 561/680 nm) before correction for spectral bleed-through, *AF*568_*em*_ is the emission in the donor channel (ex/em 561/586 nm), and *AF*647_*em*_ is the emission in the acceptor channel (ex/em 660/680 nm). The coefficient *α* corrects the bleed-through contribution from the donor (Alexa568) due to the overlap of the donor emission in the FRET channel. The coefficient *β* corrects the non-FRET bleed-through from the acceptor (Alexa647) due to the excitation of the acceptor at the donor excitation wavelength. Correction factors *α* and *β* were determined independently for each experiment (see Methods). Simultaneous with the observation of lysosome docking on phagosomes, the FRET ratio of single phagosomes increased continuously with time until it reached a plateau, following a sigmoidal relationship (Fig. 2d). This is a general phenomenon confirmed in over 40 phagosomes from 22 cells (Supplementary Fig. 15a-c). The gradual increase in FRET ratio as a function of time indicates an increasing concentration of acceptors (Alexa647) in phagosomes resulted from their continuous fusion with lysosomes. FRET signal was diminished when microtubules were disrupted by nocodazole treatment, which further confirms that this assay measures phagosome-lysosome fusion (Supplementary Fig. 16).

We then asked: what caused the plateau in the FRET ratio of single phagosomes? This is unlikely to be caused by the completion of phagosome-lysosome fusion, because lysosomes were seen to dock on phagosomes after the FRET ratio reached its plateau. We explored two possible causes for the plateau seen in the FRET ratio vs. time plots. One is that the concentration of acceptor fluorophores delivered into phagosomes by lysosome fusion reached the threshold needed to saturate the donor fluorophores on RotSensors. The second is that the acceptor concentration in the phagosome had reached equilibrium with that in the lysosomes that were fusing with it. Even as more lysosomes continued to fuse with the phagosome, the content exchange would no longer increase the concentration of acceptors in the phagosome. To determine which of these two possible circumstances caused the FRET ratio to plateau, we varied [BSA-Alexa647-biotin]_loading_ from 2.5 µg/mL to 25 µg/mL (note: concentrations higher than 25 µg/mL resulted in decreased cell viability). While the exact concentration of acceptors (Alexa647) inside lysosomes is unknown, it is proportional to [BSA-Alexa647-biotin]_loading_ (Supplementary Fig. 14 and Supplementary Fig. 15a-c). The rationale here is that if the plateau in FRET ratio was caused by the saturation of donors on RotSensors, the magnitude of plateau would be determined by the amount of donors on RotSensors and thus should not change with [BSA-Alexa647-biotin]_loading_. On the contrary, if the plateau of FRET ratio occurred because the acceptor concentration reached an equilibrium between the phagosome and fusing lysosomes, the magnitude of plateau would be determined by the acceptor concentration inside lysosomes and thus should increase with [BSA-Alexa647-biotin]_loading_. In experiments, we found that the plateau level of FRET ratio increased with [BSA-Alexa647-biotin]_loading_ from 2.5 to 25 µg/mL (Supplementary Fig. 15a-d). This indicates that the FRET ratio plateau was due to the establishment of equilibrium between the phagosome lumen and lysosomes, and not saturation of donors on RotSensors.

We further optimized [BSA-Alexa647-biotin]_loading_ to minimize cell-to-cell variation in loading BSA-Alexa647-biotin into lysosomes. We found that the higher [BSA-Alexa647-biotin]_loading_, the more variation in the amount of BSA-Alexa647-biotin loaded into lysosomes between cells (Supplementary Fig. 14). As a result, the plateau level of FRET ratio had larger variations at [BSA-Alexa647-biotin]_loading_ > 10 µg/mL (Supplementary Fig. 15d). To estimate how the cell-to-cell variation affects the measurement of phagosome-lysosome fusion kinetics, we normalized the FRET ratio vs. time plot of each phagosome to its plateau level and then from the sigmoidal Boltzmann fitting of each FRET ratio plot, we calculated the slope at half-response point *t*_0_ as the kinetic rate of phagosome-lysosome fusion (referred to as FRET rate) (Fig. 2d and Supplementary Fig. 15e). In the FRET rate vs. [BSA-Alexa647-biotin]_loading_ plot (Supplementary Fig. 15f), the standard deviation of FRET rate between phagosomes increased significantly at [BSA-Alexa647-biotin]_loading_ > 10 µg/ml. All results suggest that an optimal range of [BSA-Alexa647-biotin]_loading_ for the FRET assay is ≤ 10 µg/ml. We therefore chose to use 10 µg/ml, the highest concentration within this range, to improve the sensitivity of the FRET assay.

### FRET-RotSensors reveal correlation between phagosome-lysosome fusion kinetics and phagosomes centripetal transport

After the validation and calibration of the FRET fusion assay, we then investigated whether the kinetics of phagosome-lysosome fusion (measured as the FRET rate) correlates with the motility of phagosomes. Much like the heterogeneity in acidification, the FRET rate differed greatly between phagosomes even within the same cell (Fig. 2e and f). However, all single phagosome FRET ratio data follow a linear correlation with the translational and rotational velocities of phagosomes, with a Pearson’s coefficient of 0.75 and 0.86, respectively (Fig. 2e and f). This indicates that phagosomes that moved faster translationally and rotated more along microtubules also fused more rapidly with lysosomes.

Because lysosomes in macrophages are more concentrated in the perinuclear region (Fig. 3a)^40^, we then asked if, besides velocity, the direction of phagosome transport plays any role in their fusion with lysosomes. To identify centripetal towards cell nucleus, we analyzed the distance of each phagosome from the center of cell nucleus as a function of time (Fig. 3b). The effective transport distance of a phagosome (grey shaded areas in Fig.3b) is the distance from the time when it begins to fuse with lysosomes to the time when it reaches the nucleus boundary. The starting time of phagosome-lysosome fusion was obtained from the sigmoidal fitting of the FRET ratio vs. time plot (Fig. 2d). We also identified segments of active centripetal motion following a previously reported method (Fig. 3b)^13^. Most phagosomes moved centripetally (Supplementary Fig. 17), but not all of them exhibited active centripetal runs (Fig. 3b). As shown in Fig. 3A, phagosome #1 underwent a few segments of active centripetal runs, but phagosome #2 did not. Concurrently, phagosome #1 also fused with lysosomes more rapidly (Fig. 3c). To confirm the generality of this result, we quantified the centripetal velocity and percentage of active centripetal runs of single phagosomes. Centripetal velocity was calculated as the effective transport distance divided by time. The percentage of active centripetal run was calculated as the sum of time of all active centripetal runs divided by the total duration of the effective transport (N = 40 phagosomes from 22 cells). The results demonstrate that phagosomes with larger centripetal velocity and higher percentage of active centripetal movements fuse faster with lysosomes (Fig. 3d).

**Fig. 3.**
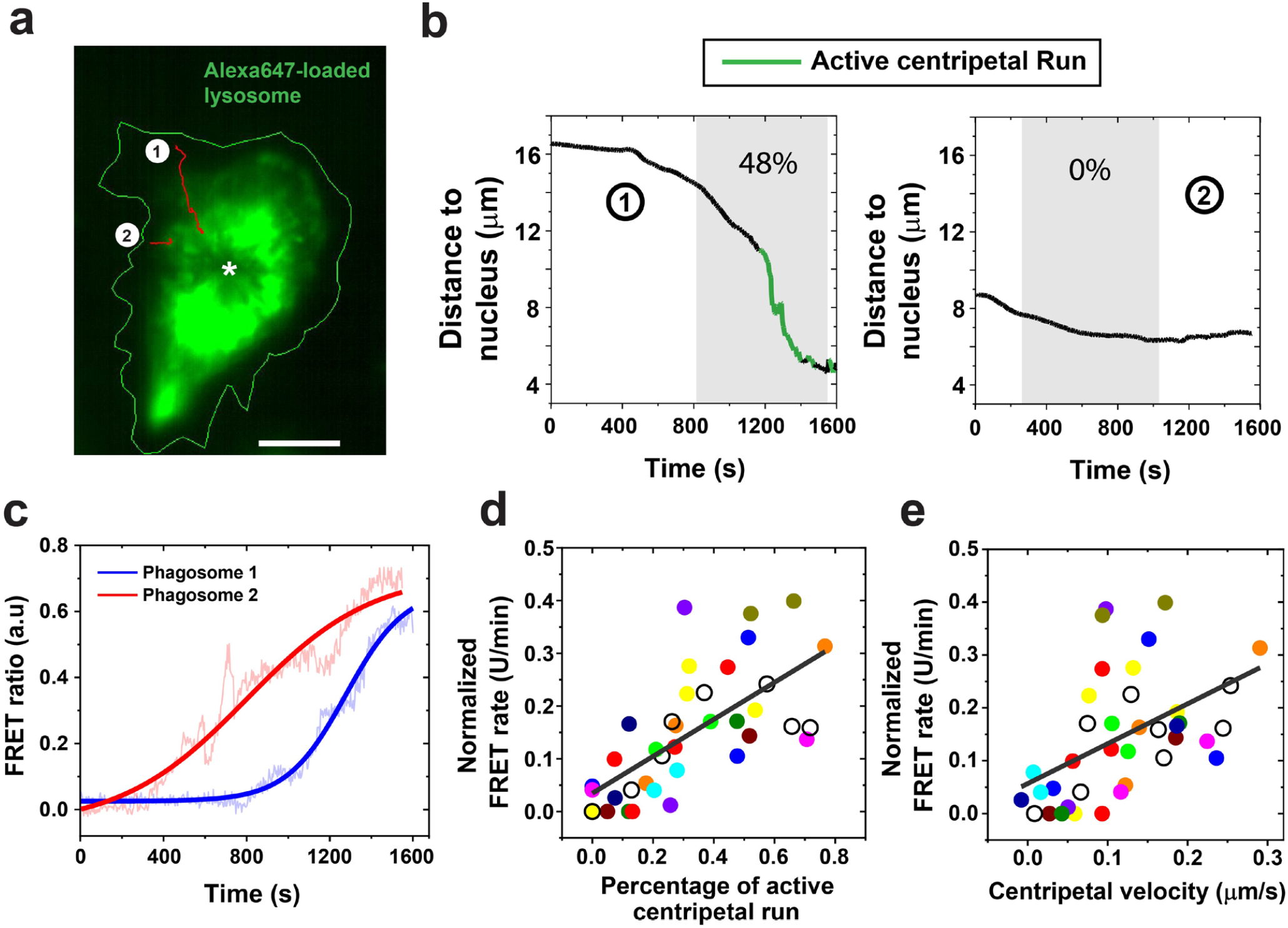
Correlation of phagosome-lysosome fusion kinetics with the centripetal motility of phagosomes. **a** Fluorescence image showing a cell in which lysosomes were labeled with BSA-Alexa647-biotin. Green line indicates the cell periphery, and the asterisk indicates the centroid of the nucleus. The initial positions of two representative phagosomes were indicated by the two number-labeled white dots and their trajectories are shown in red. Scale bar, 10 μm. **b** Plots showing the phagosome-to-nucleus distance as a function of time for both the phagosomes indicated in **a**. Segments of active centripetal runs are highlighted in green. Grey shaded area indicates the effective transport distance of the phagosome from the time where it begins to fuse with lysosomes to the time when it reaches the nucleus boundary. The starting time of phagosome lysosome fusion was obtained based on sigmoidal fitting the FRET ratio vs. time plot. Phagosome #1 underwent active centripetal runs in 48% of the time during effective transport, whereas phagosome #2 had no active centripetal run. **c** FRET ratio vs. time plots for both phagosomes marked in **a**. Solid lines indicate sigmoidal-Boltzmann fitting to the data. **d** and **e** Scatter plots showing normalized FRET rate of single phagosomes plotted against the percentage of active centripetal motion (**d**) and centripetal velocity (**e**). Each data point represents data from a single phagosome. Data from multiple phagosomes inside the same cell are shown in the same solid color. Data from cells in which only one phagosome was studied are shown as black circles. N = 40 phagosomes from 22 cells. The black lines indicate linear regression with a Pearson’s coefficient of 0.68 and 0.49, respectively.

### MagSensors for magnetic manipulation and imaging of phagosomes

To determine whether the centripetal motility of phagosomes determines their fusion with lysosomes and consequently their acidification, we applied magnetic tweezers to manipulate the intracellular transport of single phagosomes. We simultaneously measured changes in phagosome degradation (Supplementary Fig. 18). We designed 1 μm magnetically modulated phagosome sensors (MagSensors) that were also pH-responsive (Fig. 4a). The MagSensors were biotinylated and conjugated with SAv-pHrodo Red (pH indicator) and SAv-CF640 (reference), using the same procedure as for the preparation of RotSensors. The MagSensors exhibited similar pH responses (Supplementary Fig. 19). In all experiments, a solenoid tip was positioned on the opposite side of the cell from the phagosome of interest, so that the phagosome could be pulled by magnetic attraction from the cell periphery towards the center. The magnetic force exerted on individual MagSensors was calibrated by tracking the movement of particles in a polydimethylsiloxane (PDMS) base calibration medium under magnetic pulling force. The magnetic force changes with the distance between the MagSensor and the solenoid tip following theoretical predictions (Fig. 4b)^41^. On average, the solenoid tip was positioned ≈ 43 μm from the MagSensor of interest. This means that the average force exerted on a MagSensor by the tip was ≈ 21 pN at the beginning of manipulation and reached as high as ≈ 31 pN as it moved closer to the tip at the end of imaging (Supplementary Fig. 20).

**Fig. 4.**
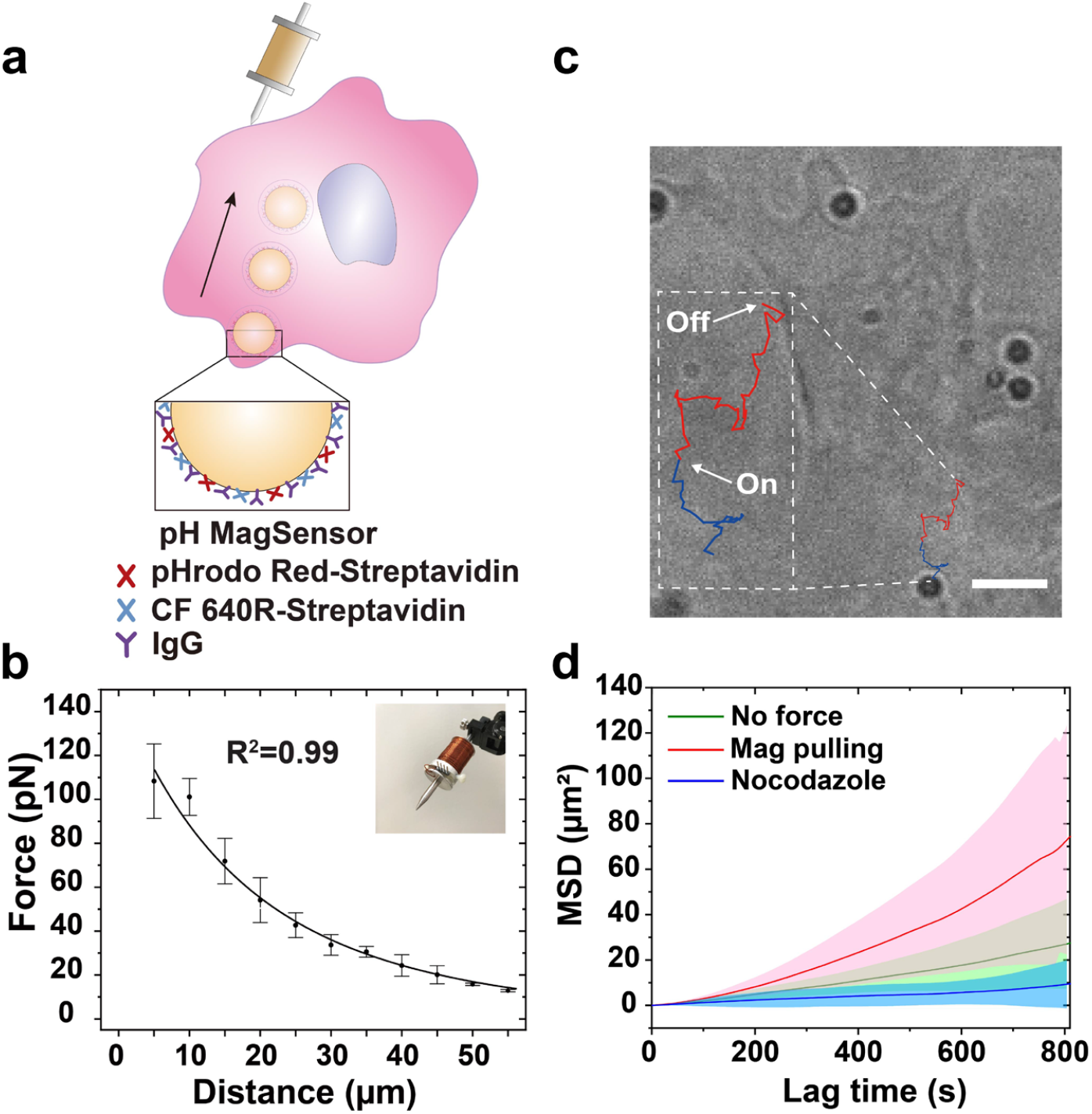
Phagosome degradation measurements during magnetically manipulated transport. **a** Schematic illustration of the experimental setup. The magnetic gradient is generated by a homemade magnetic tweezers setup built on an inverted fluorescence microscope system. Magnetic pulling force was applied on MagSensors after their internalization into phagosomes. The 1 µm MagSensors were coated with SAv-pHrodo Red (pH indicator), SAv-CF640 (reference), and physically adsorbed IgG. **b** Calibration plot showing the magnetic force exerted on each MagSensor as a function of its distance to the tip of the magnetic tweezers solenoid (shown in inset). Error bars are standard deviations from 5 samples. **c** Bright-field image of a cell overlaid with the trajectory of a MagSensor-containing phagosome under magnetic pulling. The start and end time points of the exertion of magnetic force are indicated. The blue-colored segment of the trajectory indicates the movement of the MagSensor before cell entry; the red-colored segment of the trajectory indicates the intracellular movement of a MagSensor-containing phagosome under magnetic manipulation. Scale bar, 5 µm. **d** Mean-square displacements (MSD) calculated from trajectories of individual MagSensors under different conditions as indicated. Each line is an average of results from N = 10 phagosomes from 8 cells (no force), 9 cells (magnetic pulling), and 7 cells (nocodazole). Shaded areas indicate standard deviations.

We performed the magnetic experiments in resting macrophages stably expressing actin-GFP (Supplementary Fig. 21). The fluorescence from actin-GFP allowed us to identify when phagosome formation completes^42, 43^, so that we can apply magnetic pulling force on the MagSensor-containing phagosomes only after their internalization. We used resting cells for these experiments because they are less flat, which allows for better magnetic manipulation of the phagosome. The acidification-motility correlation of phagosomes in resting cells was similar to that in activated cells (Supplementary Fig. 6). In the experiments, we turned on the magnetic force immediately after the MagSensors were internalized into phagosomes, which was indicated by the actin-GFP intensity peak, and pulled the MagSensor-containing phagosomes from the cell periphery towards the center (Fig. 4a). The magnetic force remained on throughout imaging. Under the magnetic pulling force, phagosomes moved towards the magnetic tip with accelerated translational velocity and larger mean square displacements (Fig. 4d and Supplementary Fig. 22). Their trajectories were zigzagged (Fig. 4c), likely because the magnetic force applied on the MagSensors was only slightly larger than the collective forces from the microtubule-based molecular motors. As mentioned above, the magnetic pulling force on single MagSensors was ≈ 21 to 31 pN (Supplementary Fig. 20). For comparison, microtubule-based molecular motors were shown to exert collective forces as high as ≈ 20 pN on vacuoles encapsulating 1 µm particles^44^. Therefore, the zigzagged movements of phagosomes are plausibly a result of the combined influence of the magnetic force and the forces exerted by molecular motors. As the translational velocity of phagosomes accelerated under magnetic pulling, they also acidified more rapidly (Fig. 5a). Phagosomes that were magnetically pulled acidified at an average rate of 0.79 ± 0.54 pH unit/min (N = 38). In contrast, for phagosomes not subject to magnetic pulling, an average rate of 0.48 ± 0.34 pH unit/min was observed (N = 33) (Fig. 5b). Surprisingly, magnetic manipulation had no effect on the final pH of the phagosomes, as phagosomes with or without magnetic manipulation reached an average final pH of 4.7 ± 0.4 and 4.7 ± 0.3, respectively (Fig. 5c). These results evidently demonstrate that faster transport of phagosomes causes their faster acidification.

**Fig. 5.**
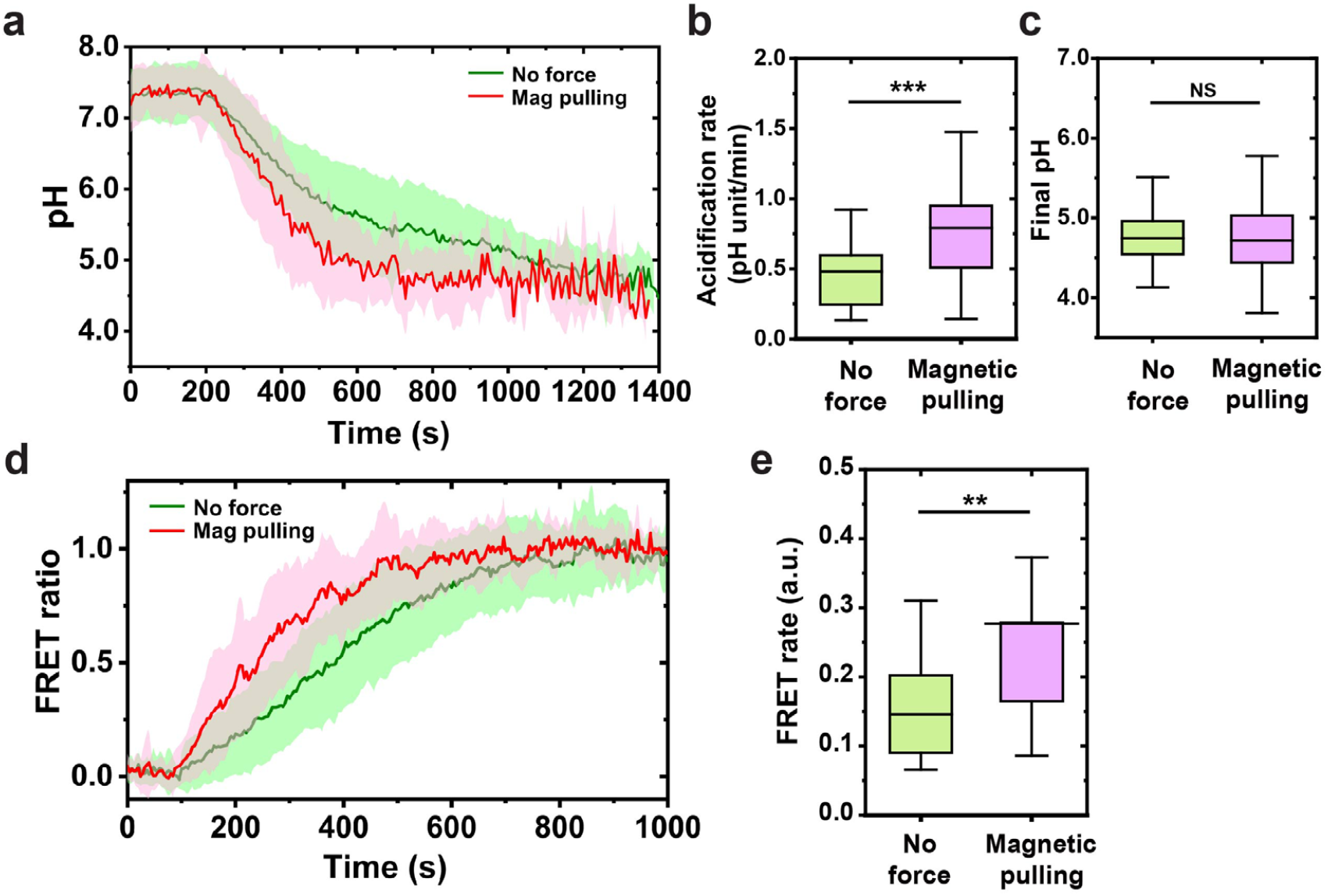
Phagosome acidification and phagosome-lysosome fusion under magnetic pulling. **a** Line plots showing the average phagosome pH as a function of time with or without magnetic pulling as indicated. Each line plot is an average from 20 phagosomes. Shaded areas represent standard deviations. **b** Box graph showing the average acidification rate of phagosomes with or without magnetic pulling. The average acidification rate is 0.48 ± 0.34 pH unit/min without magnetic pulling (N = 33 phagosomes in 29 cells from 11 independent experiments) and 0.79 ± 0.54 pH unit/min with magnetic pulling (N = 38 phagosomes in 38 cells from 13 independent experiments). **c** Box graph showing the average final pH in different experiment conditions as indicated. The average final pH is 4.7 ± 0.3 without magnetic manipulation (N = 33) and 4.7 ± 0.4 with magnetic pulling (N = 38). **d** Line plots showing the average normalized FRET ratio as a function of time with or without magnetic pulling as indicated. The line curves are averaged from 20 individual phagosomes in each experimental condition. Shaded areas represent standard deviations. **e** Box graph showing the average normalized FRET rate in different experiment conditions as indicated. The average FRET rate is 0.15 ± 0.07 without magnetic manipulation (N = 20 phagosomes in 16 cells from 4 independent experiments) and 0.28 ± 0.21 with magnetic pulling (N = 20 phagosomes in 19 cells from 8 independent experiments). In **b, c** and **e**, each box plot indicates the mean (horizontal line) and the interquartile range from 25% to 75% of the corresponding data set. Statistical significance is highlighted by p-values (Mann-Whitney Test) as follows: ** p < 0.01, *** p < 0.0001, NS p > 0.05.

We next investigated how magnetic pulling affects phagosome-lysosome fusion using the FRET fusion assay shown in Fig. 2. BSA-Alexa647-biotin (10 µg/ml) was used to load lysosomes. As shown in Fig. 5c, the FRET signals from magnetically pulled phagosomes (average from N = 20 each) were noticeably more intense and increased more rapidly, indicating enhanced phagosome fusion with lysosomes. The FRET rate, which indicates the phagosome-lysosome fusion kinetic rate, is 0.28 ± 0.21 a.u. (N = 20) for phagosomes under magnetic pulling, in contrast to the average rate of 0.15 ± 0.07 a.u. without magnetic forces (N = 20) (Fig. 5e). To investigate whether the movement of phagosomes alone is sufficient to facilitate productive fusion with lysosomes, we next treated cells with nocodazole, which disrupts microtubule-based transport of both phagosomes and lysosomes. We then magnetically moved phagosomes the same way as in non-treated cells. Despite the fast movement of phagosomes under magnetic pulling forces, they exhibited minimal fusion with lysosomes and minimal acidification (Supplementary Fig. 23). This suggests that productive phagosome-lysosome fusion requires active transport of not only the phagosomes but also the lysosomes that they fuse with.

The magnetic tweezers results, taken together, demonstrate that faster transport of phagosomes on microtubules promotes their encounters with lysosomes, which leads to faster acidification. This explains why phagosome transport velocity positively correlates with phagosome-lysosome fusion kinetics and acidification rate.

## Discussion

Phagosomes, even those from the same cells, show a significant amount of variation from one another in their degradation and transport. Each phagosome functions as a separate degradative unit with its own distinct degradation kinetics and transport velocity. However, the individuality of phagosomes has been overlooked in previous studies that relied on population-based measurements of phagosomes^8, 30, 45, 46^. There is no known quantitative relationship between the seemingly stochastic rates of degradation of single phagosomes and their equally stochastic transport dynamics. In the current study, we detail the development of a single-phagosome imaging and manipulation toolset that allowed us to probe both degradative processes and dynamics of individual phagosomes. Multimodal particle sensors, called RotSensors, were engineered as phagosome probes to measure acidification within the lumen of an individual phagosome, the kinetics of that same phagosome’s fusion with lysosomes, and its overall rotational and translational motion within the cell. The RotSensors were further combined with magnetic tweezers to perturb phagosome transport and observe the consequences for the degradation of single phagosomes. This integrated approach allowed us to demonstrate that the biochemical progression of degradative functions in individual phagosomes is coupled to and determined by the dynamics of their active transport.

By simultaneously tracking the transport dynamics and degradation of single phagosomes, we first determined that phagosomes that move faster centripetally fuse with lysosomes and acidify faster. While phagosome degradation is known to require microtubule-dependent transport^19, 21, 33, 47, 48, 49, 50, 51^, our results demonstrate that the seemingly stochastic rate of degradation of single phagosomes is positively correlated with their equally stochastic velocity of transport. In the magnetic tweezers experiments, we then showed that when phagosomes are accelerated towards nucleus, they speed up their fusion with lysosomes and their acidification. The velocity of phagosomes directly regulates where and how fast they fuse with lysosomes, and this, in turn, determines the acidification. Surprisingly however, phagosome velocity doesn’t influence the final pH of the phagosomes. Our results suggest that centripetal transport of phagosomes functions as a clock that determines the timing of their sequential biochemical activities during pathogen degradation. The precise positioning of phagosomes largely relies on microtubules-based motors^52^. It was shown previously that Rab7-RILP (Rab7-interacting lysosomal protein) complex induces the recruitment of the dynein-dynactin motor protein complex to phagosomes^53^, and the clustering of dynein generates extra force to mediate the rapid centripetal transport of phagosomes towards the perinuclear region^50^. Because lysosomes are enriched in perinuclear region (Fig. 3a) and the perinuclear lysosomes are more acidic than those near cell periphery^54, 55^, it is possible that dynein-mediated active centripetal transport enhances the probability of physical collision between a phagosome and lysosomes, hence promoting the degradation of phagosome lumen. We further confirmed that it is the active microtubule-based centripetal transport, rather than simple diffusion, regulates phagosome degradation. Because phagosomes failed to fuse with lysosomes or to acidify in cells with disrupted microtubules, even if phagosomes were moved magnetically. Moreover, Rab7 not only bridges phagosomes with dynein-dynactin, but also directly recruits the homotypic fusion and vacuole protein sorting (HOPS) complex and V-ATPase subunit V1G1, which are involved in membrane fusion and phagosome acidification^56, 57^. Those literature reports, albeit scattered, suggest that phagosome membrane associated molecules, such as Rab7-RILP complexes, may function as the molecular links that relate phagosome motility with their degradation activities.

In addition to studying phagosomes, we envision that the RotSensor toolset presented here could be further applied, with appropriate modifications, to studying intracellular pathogens and endosome degradation. Many pathogens, such as the bacterium *Legionella pneumophila*, hijack the phagosome-lysosome fusion process to evade immune clearance^9, 10, 11^. Our study here was done with synthetic beads, which are far less complex than pathogens. Nevertheless, the results imply that intracellular pathogens might prolong their survival in host cells by disrupting the intracellular transport of phagosomes to slow down fusion with lysosomes and thus delay the degradation. This hypothesis can be tested in future studies where pathogen-extracted bioparticles or intact pathogens, instead of synthetic particles, can be used in the single-particle assays. Similarly, when smaller nanoparticles are used to replace the micron-sized beads in the single-particle assays, it will be possible to study the degradative function of endosomes that are smaller than phagosomes.

## Methods Materials

Carboxylate-modified yellow-green fluorescent polystyrene nanoparticles (diameter 100 nm), carboxylate-modified superparamagnetic Dynabeads (diameter 1 µm), pentylamine-biotin, Alexa Fluor 568 NHS ester (succinimidyl ester), Alexa Fluor 647 NHS ester (succinimidyl ester), pHrodo iFL Red STP Ester, Streptavidin, LysoTracker Green DND-26, Alexa Fluor 488 anti-tubulin-α antibody, Alexa Fluor Plus 647 phalloidin, and DQ-green BSA were purchased from ThermoFisher (Waltham, MA). Immunoglobulin G from rabbit plasma, albumin from bovine serum (BSA), biotin N-hydroxysuccinimide ester (biotin-NHS), nocodazole, and lipopolysaccharides were purchased from Sigma-Aldrich (St. Louis, MO). Monodisperse amine-modified silica particles (diameter 1.0 μm) were purchased from Spherotech Inc. (Lake Forest, IL). Sylgard 184 PDMS base was from Dow Corning (Midland, MI). 1-(3-Dimethylaminopropyl)-3-ethylcarbodiimide hydrochloride (EDC) was purchased from Alfa Aesar (Haverhill, MA). Nigericin sodium salt was purchased from Tocris Bioscience (Minneapolis, MN). Recombinant Murine IFN-γ was purchased from Peprotech (Rocky Hill, NJ). CF640R-amine was purchased from Biotium (Fremont, CA). RAW264.7 macrophage was purchased from ATCC (Manassas, VA). RAW264.7 macrophages stably expressing EGFP-actin have been previously described ^42^. Ringer’s solution (pH = 7.3, 10 mM HEPES, 10 mM glucose, 155 mM NaCl, 2 mM NaH_2_PO_4_, 5 mM KCl, 2 mM CaCl_2_, 1 mM MgCl_2_) was used for live-cell imaging. Potassium-rich solution (135 mM KCl, 2 mM K_2_HPO_2,_ 1.2 mM CaCl_2_, 0.8 mM MgSO_4_) was used for intracellular pH calibration. Acidic washing solution (135 mM KCl, 2 mM K_2_HPO_2,_ 1.2 mM CaCl_2_, 0.8 mM MgSO_4,_ 5 mM sodium citrate) at pH of 4.5 was used for particle washing in the EDC coupling step of the RotSensor fabrication. Artificial lysosome fluid (55 mM NaCl, 0.5 mM Na_2_HPO_4_, 0.26 mM trisodium citrate dihydrate, 0.79 mM glycine, 150 mM NaOH, 108 mM citric acid, 0.87 mM CaCl_2_·2H_2_O, 0.27 mM Na_2_SO_4_, 0.25 mM MgCl_2_·6H_2_O, 0.46 mM disodium tartrate, 1.6 mM sodium pyruvate) was prepared following a previously reported protocol ^58^ and used for washing particles during the protein conjugation step of the RotSensor fabrication.

### Cell Culture, Pharmacological Treatments

Both RAW264.7 macrophage and stable cell line expressing EGFP-actin were cultured in Dulbecco’s Modified Eagle Medium (DMEM) complete medium supplemented with 10% fetal bovine serum (FBS), 100 units/ml penicillin and 100 µg/ml streptomycin at 37°C and 5% CO_2._ Resting macrophages were activated with a combination of 50 ng/ml LPS and 100 units/ml IFN-γ for 9 h prior to live cell imaging. Microtubule depolymerization was achieved by incubating cells with 10 µM nocodazole for 1 h prior to imaging, and 10 µM nocodazole was presented during live cell imaging.

### Fabrication and Characterization of Phagosome Sensors

#### (a) Fluorescence labeling of Streptavidin and Bovine Serum Albumin (BSA)

To prepare streptavidin-pHrodo Red conjugates (SAv-pHrodo Red), 0.7 mg of streptavidin, 60 µg of pHrodo Red STP Ester were mixed in 350 µl NaHCO_3_ solution (100mM, pH 8.2) for 3 h at room temperature. Free dyes were removed by centrifugal filtration using Amicon Ultra filters (30K). To prepare streptavidin-CF640R conjugates (SAv-CF640), 0.7 mg of streptavidin, 80 µg of CF640R-amine, and 3 mg of EDC were mixed in 350 µl MES buffer (50mM, pH 4.5) for 3 h at room temperature. Free dyes were removed by centrifugal filtration using Amicon Ultra filters (30K). To prepare streptavidin-Alexa568 conjugates (SAv-Alexa568), 0.5 mg of streptavidin, 60 µg of Alexa Fluor 568 NHS ester were mixed in 235 µl NaHCO_3_ solution (100mM, pH 8.2) for 1 h at room temperature. Free dyes were removed by centrifugal filtration using Amicon Ultra filters (30K). To prepare bovine serum albumin-Alexa647-biotin (BSA-Alexa647-biotin) conjugates, a mixture containing 2 mg/ml BSA and 188 µg/ml (molar ratio of 1: 20) biotin-NHS in NaHCO_3_ solution (100 mM, pH 8.2) was incubated at room temperature for 1 h. Unbound biotin-NHS was removed by centrifugal filtration using Amicon Ultra filters (30K). After washing, a mixture containing 2 mg/ml BSA-biotin and 780 µg/ml Alexa Fluor 647 NHS ester in NaHCO_3_ solution (100 mM, pH 8.2) was incubated at room temperature for 3 h. Free dyes were removed by centrifugal filtration using Amicon Ultra filters (30K). The resulting BSA-Alexa647-biotin yielded dye: protein ratios of 1.2:1 on NanoDrop UV-Vis measurement (ThermoFisher, Waltham, MA). To prepare bovine serum albumin-Alexa647 (BSA-Alexa647) conjugates, a mixture containing 2 mg/ml BSA and 780 µg/ml Alexa Fluor 647 NHS ester in NaHCO_3_ solution (100 mM, pH 8.2) was incubated at room temperature for 1 h. Free dyes were removed by centrifugal filtration using Amicon Ultra filters (30K). The resulting BSA-Alexa647 yielded dye: protein ratios of 1.3:1 based on NanoDrop UV-Vis measurement (ThermoFisher, Waltham, MA).

#### (b) Fabrication of Rotational Phagosome Sensors (RotSensors)

RotSensors: Amine-modified non-fluorescent silica particles (1 µm) were incubated with 150 µg/ml biotin-NHS in NaHCO_3_ solution (10 mM, pH 8.25) for 2 h at room temperature and then with 500 µg/ml of biotin-NHS for 30 min for a second round of biotinylation. After biotinylation, silica particles were washed in methanol and deionized (DI) water. The “snowman”-like design was introduced by covalently conjugating the carboxylate-modified yellow-green fluorescent nanoparticles (100 nm) to the biotinylated and amine-modified silica particles (1 µm) using EDC coupling. In brief, carboxylate-modified yellow-green fluorescent nanoparticles (100 nm) were mixed with the biotinylated and amine-modified silica particles (1 µm) at a molar ratio of 333:1 in phosphate buffer (10 mM, pH 7.0) containing 1.0 mg/ml EDC for 2 h at room temperature. After the EDC coupling, particles were washed with acidic washing solution and 1×PBS to remove the non-covalently bounded carboxylate-modified yellow-green fluorescent nanoparticles (100 nm). 20% of the particles were found to have the “snowman” shape with 1:1 coupling ratio of the biotinylated and amine-modified silica particles (1 µm) and the carboxylate-modified yellow-green fluorescent nanoparticles (100 nm).

pH-RotSensors: pH-RotSensors were fabricated by labeling RotSensors with SAv-pHrodo Red and SAv-CF640. RotSensors were incubated with 25 µg/ml of SAv-pHrodo Red, 2.5 µg/ml of SAv-CF640, 5 µg/ml of BSA and 1 µg/ml of IgG in 1×PBS for 5 h at room temperature. Unbound proteins were rinsed off with artificial lysosome fluid and 1×PBS. The resulting pH-RotSensors were further opsonized with 30 µg/ml IgG in 1×PBS for additional 2 h before adding to cell samples.

FRET-RotSensors: FRET-RotSensors were fabricated by labeling RotSensors with SAv-Alexa 568. The RotSensors were incubated with 27.5 µg/ml of SAv-Alexa568, 5 µg/ml of BSA, and 1 µg/ml of IgG in 1×PBS for 5 h at room temperature. Unbound proteins were rinsed off with artificial lysosome solution and 1×PBS. The FRET-RotSensors were further opsonized with 30 µg/ml of IgG in 1×PBS for additional 2 h before adding to cell samples.

#### (c) Fabrication of Magnetically Modulated Phagosome Sensors (MagSensors)

pH-MagSensors: The first step is the biotinylation of carboxylate-modified Dynabeads (1 µm). In brief, 10 µl of stock Dynabeads suspension (10 mg/ml) were added to 100 µl of MES buffer (50 mM, pH 6.2) containing 10 mg/ml EDC and 1 mM of biotin pentylamine. After 1-h incubation at room temperature, biotinylated Dynabeads (1 µm) were washed in 1×PBS to remove unbound biotin pentylamine. pH-MagSensors were fabricated by labeling biotinylated Dynabeads (1 µm) with SAv-pHrodo Red and SAv-CF640. Biotinylated Dynabeads were incubated with 50 µg/ml of SAv-pHrodo Red and 50 µg/ml of SAv-CF640 for 1 h at room temperature. The resulting pH-MagSensors were further opsonized with 1 mg/ml of IgG in 1×PBS for additional 1 h before adding to cell samples.

FRET-MagSensors: Biotinylated Dynabeads (1 µm) were prepared following the same procedure described above. FRET-MagSensors were fabricated by labeling biotinylated Dynabeads (1 µm) with SAv-Alexa568. Biotinylated Dynabeads were incubated with 27.5 µg/ml of SAv-Alexa568 for 1 h at room temperature. The resulting FRET-MagSensors were further opsonized with 1 mg/ml of IgG in 1×PBS for additional 1 h before adding to cell samples.

### Magnetic Tweezers Setup and Force Calibration

Magnetic tweezers were built on an inverted fluorescence microscope system (Nikon Eclipse Ti-U, Nikon, Tokyo, Japan), as shown in Supplementary Fig. 18. The setup mainly includes a solenoid and a power supply. The solenoid was assembled by inserting a high permeability HyMu-80 alloy rod (Carpenter Technology, Reading, PA) into an aluminum bobbin wrapped with 600 turns of copper coil ^59^ (Supplementary Fig. 18b). The tip of the rod has a diameter of ≈ 1 µm (Supplementary Fig. 18c). Position of the solenoid was controlled with a manual micromanipulator (Narishige NMN-21) to achieve independent control of its position in the x-, y-, and z-direction with a minimum graduation of 250 nm in the x-y plane and 1 µm in the z-direction. The current going through the solenoid was generated by a programmable power supply (Tekpower, Montclair, CA) with a maximum power output of 5 A.

Magnetic force was calibrated as a function of particle-to-tip distance by measuring the movements of Dynabeads (1 µm) in magnetic field. In brief, CF640R labeled Dynabeads (1 µm) were suspended in PDMS base at a concentration of ≈ 3.0 × 10^5^ particles/ml. This low particle concentration was necessary to avoid particle aggregation and inter-particle magnetic inference ^60^. The PDMS base, which has a high viscosity of 5.1 Pa·s, was chosen to slow down particle movements to make particle tracking feasible. The initial positions of the solenoid tip and of the magnetic particles were imaged in bright field before time-lapse epi-fluorescence images of particles were acquired with an interval time of 0.2 s. Working current of the magnetic tweezers was 1.0 A in the calibration. The magnetic force *F*(*r*) exerted on each magnetic particles at particle-to-tip distance r was converted from particle velocity values using Stokes-Einstein equation:

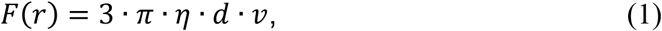

where *η* is the viscosity of calibration solution PDMS base, d represents the diameter of particle, and v is the velocity of particle. The relationship between magnetic force F(r) and particle-to-tip distance *r* was fitted using equation:

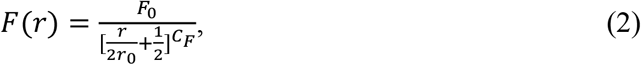

Where *F*_0_ is a force constant with the unit of pN, *r*_0_ is a distance constant with the unit of µm, *C*_*F*_ is unitless and *F*_(*p*0)_ = *F*_0_ ^59^.

### Fluorescence Microscopy

#### (a) Single-phagosome pH Assay

RAW264.7 macrophage cells were seeded on glass coverslips at 2.0 × 10^5^ cells/ml in complete medium for 24 h and then activated with LPS and IFN-γ following the procedure described above. For all single-phagosome pH studies, the recognition of pH-Sensors (i.e. pH-RotSensor and pH-MagSensor) by macrophages was synchronized following a previously reported protocol ^61^. Briefly, cell samples were cooled on ice for 3 min to postpone phagocytosis. pH-Sensors were then added at a particle-to-cell ratio of ∼5:1. Bindings of particles on macrophage were synchronized by centrifuging cell samples at 200×g for 30 s. Live cell imaging was conducted in Ringer’s solution at 37°C on a Nikon Eclipse-Ti inverted microscope equipped with a 1.49 N.A. ×100 TIRF objective (Nikon, Tokyo, Japan) and an Andor iXon3 EMCCD camera (Andor Technology, Belfast, U.K.). Fluorescence emissions at three wavelengths (ex: 488, 561, and 660 nm; em: 515, 586 and 680 nm) were acquired for the time-lapse imaging. The acquisition rate was 2 sec/frame. Single-phagosome pH assay in actin-GFP expressing RAW264.7 macrophages was conducted following the same procedure described above.

#### (b) Single-phagosome FRET-fusion Assay

RAW264.7 macrophage cells were seeded on glass coverslips at 2.0 × 10^5^ cells/ml in complete medium for 24 h and then activated with LPS and IFN-γ following the procedure described above. Loading of either BSA-Alexa647-biotin or BSA-Alexa647 into lysosome compartment was carried out following a previously reported pulse-chase protocol ^62, 63, 64^. In brief, RAW264.7 macrophage cells were incubated overnight in complete medium containing either BSA-Alexa647-biotin or BSA-Alexa647 at indicated concentration. 2 h prior to live cell imaging, cells were rinsed twice with complete medium. Labeled endocytic compartment was then chased at 37°C for 2 h to ensure complete accumulation and fragmentation of fluorescently labeled BSA in lysosome compartments (Supplementary Fig. 11 and Supplementary Fig. 12). Synchronized internalization of FRET-Sensors (i.e. FRET-RotSensor and FRET-MagSensor) was conducted following the same procedure described above. Time-lapse epi-fluorescence images were acquired to record FRET emission (FRET_0_; ex/em 561/680 nm), donor emission (Alexa568_em_; ex/em 561/586 nm), acceptor emission (Alexa647_em_; ex/em 660/680 nm) and the emission from 100 nm yellow-green fluorescence particle (ex/em 488/515 nm). The acquisition rate was 4 sec/frame.

The broad emission spectra of Alexa568 (donor) overlap partially with the acceptor emission channel (680 nm) and the broad excitation spectra of Alexa647 (acceptor) did overlap partially with donor excitation channel (561 nm). This resulted in two types of spectral cross-talks: (i) the detection of donor fluorophore (Alexa568) emission at acceptor emission channel (680 nm) under the donor excitation of 561 nm and (ii) the detection of acceptor fluorophore (Alexa647) emission at acceptor emission channel (680 nm) under the donor excitation of 561 nm in the absence of FRET effect. These two cross-talks, however, are constant throughout FRET-fusion assay, and could be deducted from the measured FRET emission (FRET_0_) to obtain FRET-generated emission (*FTET*_*em*_). Correction factors *α* and *β* for cross-talks were determined for each independent experiment based on: (1) the measured FRET emission and donor emission of Alexa568 streptavidin particles outside cells and (2) the measured FRET emission and acceptor emission of acceptor fluorophore (Alexa647) in cells without FRET-sensor. The FRET ratio is obtained as follows:

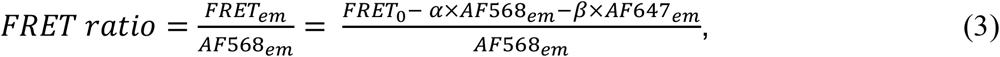

where *FTET*_0_ is the measured FRET emission (ex/em 561/680 nm) before cross-talk correction, *FTET*_*em*_ is the FRET emission, *AF*568_*em*_ is the donor emission (ex/em 561/586 nm) and *AF*647_*em*_ is the acceptor emission (ex/em 660/680 nm). The FRET increase was gradual, not stepwise, because multiple dye-loaded lysosomes can simultaneously fuse with a single phagosome and the increment in FRET signal from a single phagosome-lysosome fusion is too small to be resolved.

#### (c) DQ-green BSA Digestion Assay

To estimate the fragmentation level of BSA-Alexa647-biotin chasing, we loaded DQ-green BSA into lysosome compartment following the same pulse-chase produced described above, and measured the dequenching kinetics of lysosomal DQ-green BSA as a function of time. A DQ-green BSA concentration at 50 µg/ml was used during pulse. After the overnight pulse, cells were rinsed 3 times with flour-free Ringer’s solution and imaged using Epi-fluorescence microscope. The emission of DQ-green (ex/em 488/515 nm) was recorded every 5 min for 2 h.

#### (d) Immunofluorescence Staining and Imaging

RAW 264.7 macrophages were seeded and activated as described above. For immunostaining of α-tubulin and F-actin, cells were first washed with 1× PBS and fixed with 2% PFA at room temperature for 5 min. Next, they were permeabilized with 0.1% Triton X-100 for 5 min at room temperature, and blocked with 2% BSA for 1 h at room temperature. After the BSA blocking, cells were incubated with 2 µg/mL Alexa Fluor 488 anti-tubulin-α antibody and 2 µg/mL Alexa Fluor 647 phalloidin for 1 h at room temperature. Super-resolution structured illumination microscopy (SIM) images of the labeled cells were acquired using a DeltaVision OMX SR imaging system equipped with a Olympus Plan Apo 60×/1.42 PSF objective and a sCMOS camera.

#### (e) Lysotracker Labeling and Imaging

RAW 264.7 macrophages were seeded and activated as described above. After overnight incubation in complete medium containing 5 µg/mL BSA-Alexa647-biotin, the labeled endocytic compartments were chased at 37°C for 1.5 h. Cells were then incubated with 50 nM Lysotracker Green DND-26 at 37°C for 30 min. After that, cells were rinsed twice with Ringer’s solution to remove the free fluorophore and imaged using Re-scan Confocal Microscopy (RCM) equipped with a 1.49 N.A. ×100 TIRF objective and ORCA-fusion CMOS camera.

### Image Analysis

#### (a) Image Registration

Image registration was done to correct for the optical shift between different imaging channels. 100 nm TetraSpeck fluorescent particles (Ex/Em: 360/430 nm, 505/515 nm, 560/580 nm, and 660/680 nm) were adsorbed on poly-l-lysine-coated glass coverslips at a surface density of ≈ 0.10 bead/μm^2^ and used as markers for image registration. After sequential imaging of the marker particles in three channels (Ex: 488, 561, and 660 nm), an affine transformation was applied to align particle localization maps from 488 and 660 nm channels (termed target images) to that from the 561 nm channel (reference image) using ImageJ plugin MultistackReg ^65^. A global mapping matrix was obtained to record all the transformation steps and used to apply the same operations to all images.

#### (b) Single-Particle Localization and Intensity Determination

The centroids of single particles in epi-fluorescence images were localized using a Gaussian-based localization algorithm in the ImageJ plugin Trackmate. To measure the emission of CF640R and pHrodo Red on pH-Sensors (i.e. pH-RotSensor and pH-MagSensor), pixel intensities within a diameter of 2 μm from the localized centroid of the particle were integrated and background-corrected using custom MATLAB algorithms. Same procedure was carried out for determining *FTET*_0_, *AF*568_*em*_ and *AF*647_*em*_ in single-phagosome FRET-fusion assay. To determine the localization uncertainties, pH-RotSensors were immobilized on poly-l-lysine-coated glass coverslip and imaged for 200 consecutive frames in Ringer’s solution. Localization uncertainty was defined as the standard deviation of the tracked particle positions in x- and y-coordinates. The localization uncertainties of the 1 µm pHrodo Red-coated particle and 100 nm yellow-green fluorescence nanoparticle were determined to be 20.6 ± 4.05 nm and 13.1 ± 2.32 nm, respectively (Supplementary Fig. 25).

#### (c) Translational and Rotational Tracking Analysis of RotSensors

The single particle tracking analysis was done as described previously ^24, 66^. Briefly, translational velocity was determined from the centroid location of the 1 µm pHrodo Red- or SAv-Alexa568-coated particle (X_red,_Y_red_) as a function of time. For rotational tracking, a vector was drawn from the centroid of the 1 µm red fluorescent particle (X_red,_Y_red_) to that of the 100 nm yellow-green nanoparticle (X_green,_Y_green_) belonging to the same RotSensor. Orientation of the vector was obtained as the in-plane angle, *φ*, for the single RotSensor. The length of the vector was used to calculate the out-of-plane angle, *θ*, using the equation below:

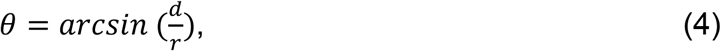

where *d* is the projection inter-particle distance between a pair of green and red dots in each given image, and *r* is the physical inter-particle distance. Because *r* varies slightly from one particle to another due to the size distribution, *r* was obtained as the maximal projected inter-particle distance *d*_*max*_ when a RotSensor samples all possible orientations and the *d*_*max*_ was larger than 550 nm. In cases when d_max_ < 550 nm, *r* was set to 550 nm. Two-dimensional rotational velocity was determined from the rotation matrix derived from *φ* and *θ* at consecutive frames. Using this method, φ can be measured within a full range (−180° to +180°), but there is ambiguity in measuring θ, because RotSensors oriented at a polar angle of θ or (180° − θ) are indistinguishable from one another when observed as a two-dimensional projection image. However, this ambiguity does not affect the measurements of Δθ and translational velocity in this study, because Δθ of phagosomes between consecutive image frames (2s/frame) is expected to be < 90°, based on the result that phagosomes rotate an average Δφ of 0.12 ± 0.16 rad (6.9° ± 9.2°) between consecutive frames (Supplementary Fig. 26). Therefore, only Δθ < 90° was used in the tracking analysis for calculating rotational velocity of phagosomes.

#### (d) Determination of Active Centripetal Run

The directional transport of phagosomes was analyzed following a previously reported method with slight modification ^13^. The centroid of the nucleus was obtained based on bright field image of the cell (Supplementary Fig. 27) ^48, 67^. At each time point of a particle trajectory, the distances between the phagosome and the centroid of the nucleus was quantified (referred to as ‘distance-to-nucleus’) (Fig. 4b). The travelled distance of phagosome towards the nucleus was subdivided into segments with a length of 32 seconds. Each segment was categorized as either a segment of active run or as a segment of passive motion based on the travelled distance with regards to nucleus. The distance threshold for active run was set to 0.5 µm, independent of the direction of motion whether it is towards the nucleus (centripetal) or towards the periphery (centrifugal) ^68^. Finally, segments of active runs were classified either as active centripetal run or as active centrifugal run depending on the directionality of the transport.

#### (e) Intracellular pH Calibration

The intracellular pH calibration of pH-RotSensors and pH-MagSensors was done following a previously published protocol ^26^. Macrophages seeded on glass coverslips were pretreated with 10 µM concanamycin in Ringer’s solution for 10 min before the addition of particles and then incubated at 37°C for 10 min to promote particle phagocytosis. Cell medium was replaced with potassium-rich buffers at different pH values. All buffers contain 20 µM of nigericin, but different buffering agent: 5 mM acetic acid for pH 4.5, 5 mM MES for pH 5.5, and 5mM HEPES for pH 6.5 and 7.3. The pH calibration was done from pH 7.3 to 4.5. For each pH, cells were rinsed twice with the correspondingly buffer and allowed to equilibrate for 5-10 min before image acquisition. Fluorescence emission of the pHrodo Red at 586 nm (*I*_*pHrodored*_) and reference dye CF640R at 680 nm (*I*_*ref*_) was obtained at various pH and background-corrected to obtain ratiometric pH calibration plots (Supplementary Fig. 2).

In live cell imaging, pH calibration was done for individual internalized pH-RotSensors and pH-MagSensors to eliminate the effect of particle-to-particle variation in their pH responses. By plotting *I*_*pHrodo*_*/I*_*ref*_ vs. time for single phagosomes, we notice that acidification process started with a standby period during which *I*_*pHrodo*_*/I*_*ref*_ ratio remains unchanged denoting that the phagosome pH remained equivalent to extracellular pH of 7.3 (Ringer’s solution, pH = 7.3) before the initiation of phagosome acidification (Supplementary Fig. 28a and b). We further confirmed this by imaging phagosome acidification in cells expressing actin-GFP, in which intensity peak of actin-GFP pinpointed the time of particle internalization ^42, 43^ (Supplementary Fig. 28c-e). After live cell imaging, the same cell sample was incubated in pH 4.5 potassium-rich pH calibration buffer containing 20 µM of nigericin to obtain the ratiometric emission (*I*_*pHrodo*_ /*I*_*ref*_) of the RotSensor at pH 4.5 (Supplementary Fig. 28a). In the final step of the calibration procedure, a linear function was generated based on the known *I*_*pHrodo*_ /*I*_*ref*_ ratios at pH 4.5 and 7.3, and the result function was used to transform the fluorescence measurements of the RotSensor to luminal pH values (Supplementary Fig. 28b and f).

#### (f) Wavelet Transform Analysis

A wavelet analysis algorithm reported by Chen *et al*. was used to distinguish active rotation from passive one. Both in-plane (*θ*) and out-of-plane (*φ*) displacement of a RotSensor was convoluted using the 1-dimensional Haar continuous time wavelet transform ^32^. The wavelet coefficients after the transformation were plotted as a function of time and for different widths of the wavelet function (Supplementary Fig. 8). The width of the wavelet function used here is the number of image frames and referred to as “scale”. Scale 10 was chosen as a universal threshold such that no active rotation of phagosomes was detected in cells that were treated with microtubule-disrupting drug nocodazole.

#### (g) Determination of location of cell entry

The boundary of the cell was manually traced in ImageJ according to the fluorescence emission (ex/em 561/586 nm) of the cell in Ringer’s solution containing trypan blue (Supplementary Fig. 27a). The boundary and the center of the nucleus was manually traced in ImageJ based on the bright field image of the given cell ^48, 69^ (Supplementary Fig. 27b).

#### (h) Calculation of Mean-Square Displacements (MSD)

Time-resolved MSD shown in Fig. 4d was analyzed using custom MATLAB algorithms. Briefly, the MSD values of each single phagosome trajectories were calculated as a function of lag time **δt** based on the following equation:

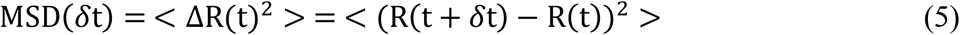

Where (R(t + *δ*t) – R(t)) is end-to-end displacement of an individual particle from the time point t to t + *δ*t.

### Transmission Electron Microscopy

Cell samples for TEM imaging were prepared based on a previously reported protocol with slight modification ^66^. Particle sensors were incubated with cells for 50 min at 37 °C before trypsinization. After centrifugation, the cell pellet was fixed on ice for 1 h by using a mixture of 2.5% (w/v) glutaraldehyde and 1% (w/v) osmium tetroxide in 1×PBS buffer. Subsequently, the pellet was stained in 0.5% (w/v) uranyl acetate aqueous solution for 12 h on ice. Cell pellets were dehydrated sequentially in a series of ice-cold aqueous solutions containing 30% (v/v), 50%, 75%, 90%, 95% and 100% ethanol for 5 min each. Dehydration in 100% (v/v) ethanol was repeated three times at room temperature. The dehydrated cell pellet was immersed sequentially in resin infiltration solutions that contain ethanol and Spurr’s resin at various volume ratio (ethanol: resin, 2:1, 1:1 and 1:2) for 30 min each at room temperature. Cells were cured in 100% resin for 18 h at 65°C prior to microtome sectioning. Sections were stained with 2% (w/v) uranyl acetate for 10 min before imaging. Samples were imaged with a JEOL JEM-1010 Transmission Electron Microscope (Electron Microscopy Center, Indiana University).

## Acknowledgments

We thank Prof. Sergio Grinstein at University of Toronto for his helpful discussion. We thank Mr. Andy Alexander at the Electronic Instrument Services of Indiana University for assistance with the setup of magnetic tweezers. We thank Dr. Barry Stein at the IUB Electron Microscopy Imaging Center, Dr. Jim Power at the IUB Light Microscopy Imaging Center (LMIC), Dr. Yi Yi at the Nanoscale Characterization Center for assistance with instrument use.

## Author contributions

Y.-q. Y., Z. Z., and Y. Y. designed research; Y.-q. Y. and Z. Z. performed research and analyzed data; G. F. W. W. contributed critical reagents; Y.-q. Y., Z. Z., and Y. Y. wrote the paper and G.F.W. W. provided critical discussion.

## Data and materials availability

All data needed to evaluate the conclusions in the paper are present in the paper and/or the Supplementary Materials. Additional data related to this paper may be requested from the

