## supplemental file for "Heterogeneous but not random: Cargo degradation in phagosomes is kinetically coupled to their intracellular mobility"

**This PDF file includes:**

Figs. S1 to S28

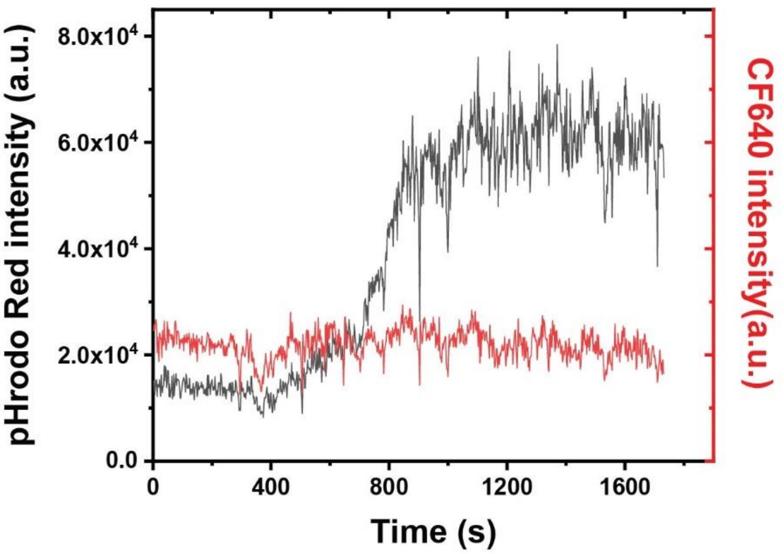


**Supplementary Fig. 1.** Line plots showing the fluorescence emission intensity vs. time of pHrodo Red and CF640R on a RotSensor during phagosome acidification. Data are representative of 57 phagosomes in 39 cells.

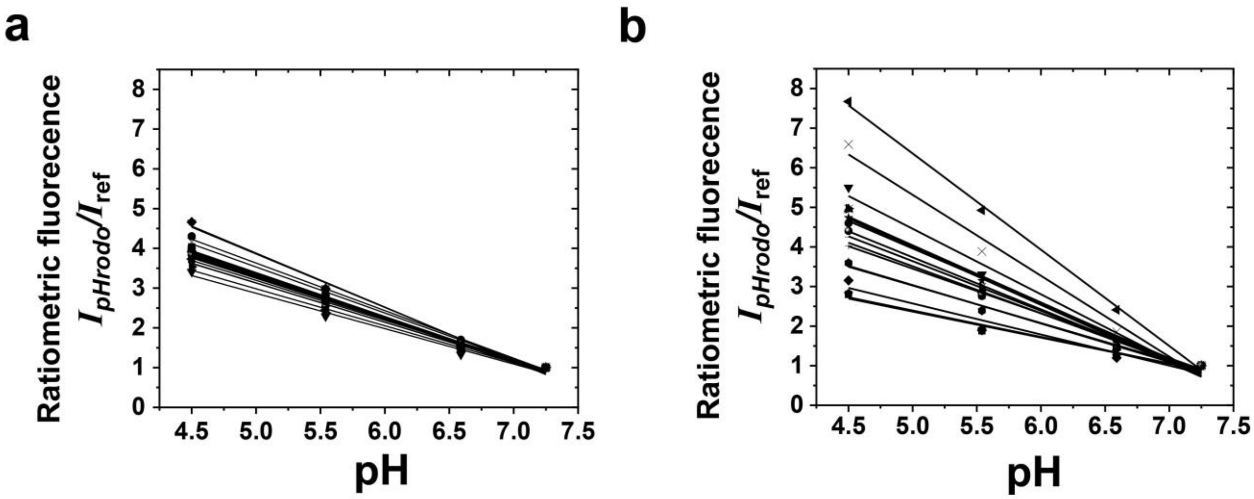


**Supplementary Fig. 2.** Extracellular (**a**) and intracellular (**b**) pH calibration plots for individual pH-RotSensors. The fluorescence emission ratio $I_{pHrodo}/I_{ref}$ at each pH was normalized to that at pH 7.3. Black lines indicate linear fittings of the data. The individual linear fittings in (**a**) (N = 18) have an average $R^2$ of 0.99 ± 0.004, and the fittings in (**b**) (N = 16) have an average $R^2$ of 0.98 ± 0.02.

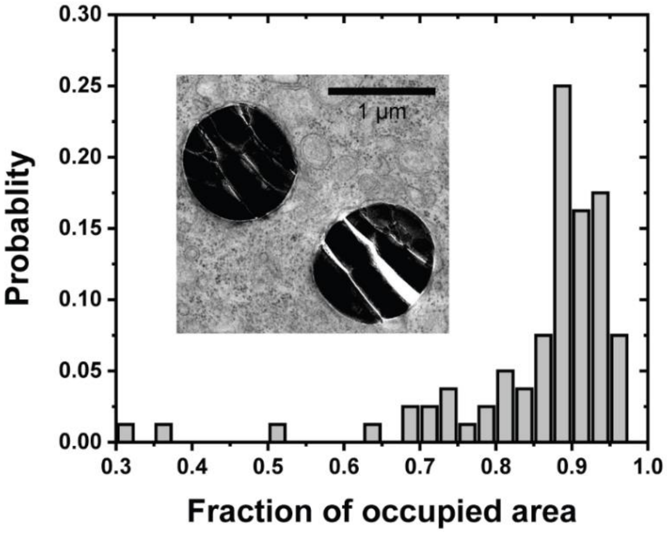


**Supplementary Fig. 3.** Characterization of the tightness of phagosome membrane encapsulating internalized particle sensors. Inset is a representative TEM image showing RotSensors encapsulated inside phagosomes. Fraction of occupied area is defined as the ratio of the area occupied by the particle sensor to the total area of the phagosome in each given TEM section. N = 80 phagosomes.

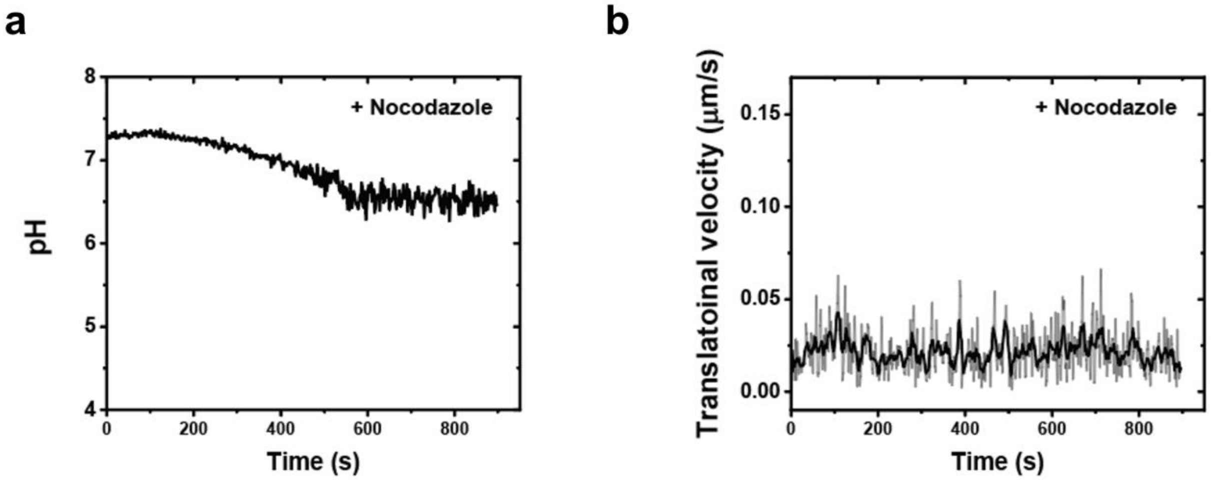


**Supplementary Fig. 4.** Acidification (**a**) and translational velocity (**b**) of a phagosome in a nocodazole-treated cell. Thick line in (**b**) indicates data after wavelet denoising. The results are representative of 8 phagosomes.

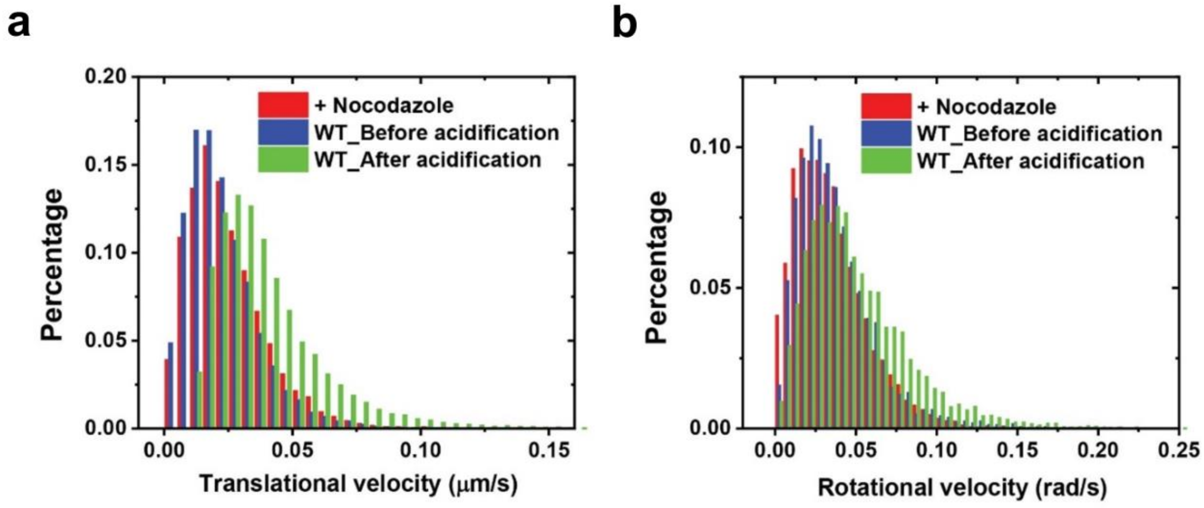


**Supplementary Fig. 5.** Histograms showing the distribution of translational (**a**) and rotational (**b**) velocities of phagosomes in control cells (WT) and cells treated with nocodazole. Data were obtained from N = 30 phagosomes from 20 control cells and N = 8 phagosomes from 8 nocodazole-treated cells.

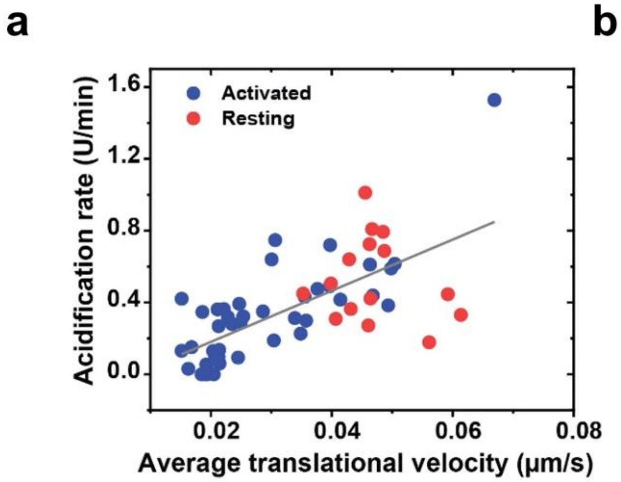


**Supplementary Fig. 6.** Scatter plots showing acidification rate against average translational (**a**) and rotational (**b**) velocities of single phagosomes during their rapid acidification period in activated and resting macrophages. Black lines indicate linear fitting to the data. Pearson coefficients are 0.68 and 0.69 for the fitting in (**a**) and (**b**), respectively. For translational tracking, N = 42 phagosomes from 24 activated cells and N = 15 phagosomes from 15 resting cells. For rotational tracking, N = 17 phagosomes from 12 activated cells and N =15 phagosomes from 15 resting cells.

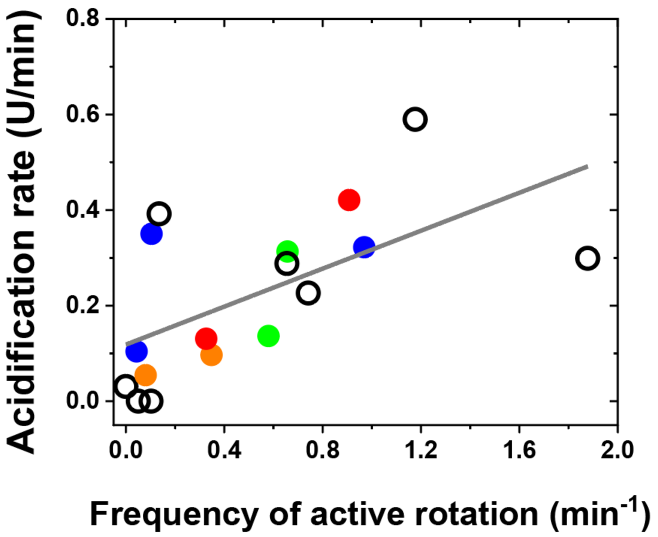


**Supplementary Fig. 7.** Scatter plot showing acidification rate against frequency of active rotation of single phagosomes during rapid acidification period. Data points from multiple phagosomes within the same cells are shown in the same solid color. Data points from cells containing only one phagosome are shown as black circles. Correlation between the phagosome acidification rate and transport velocities is indicated by the linear regression with a Pearson's coefficient of 0.60. N = 17 phagosomes from 12 cells.

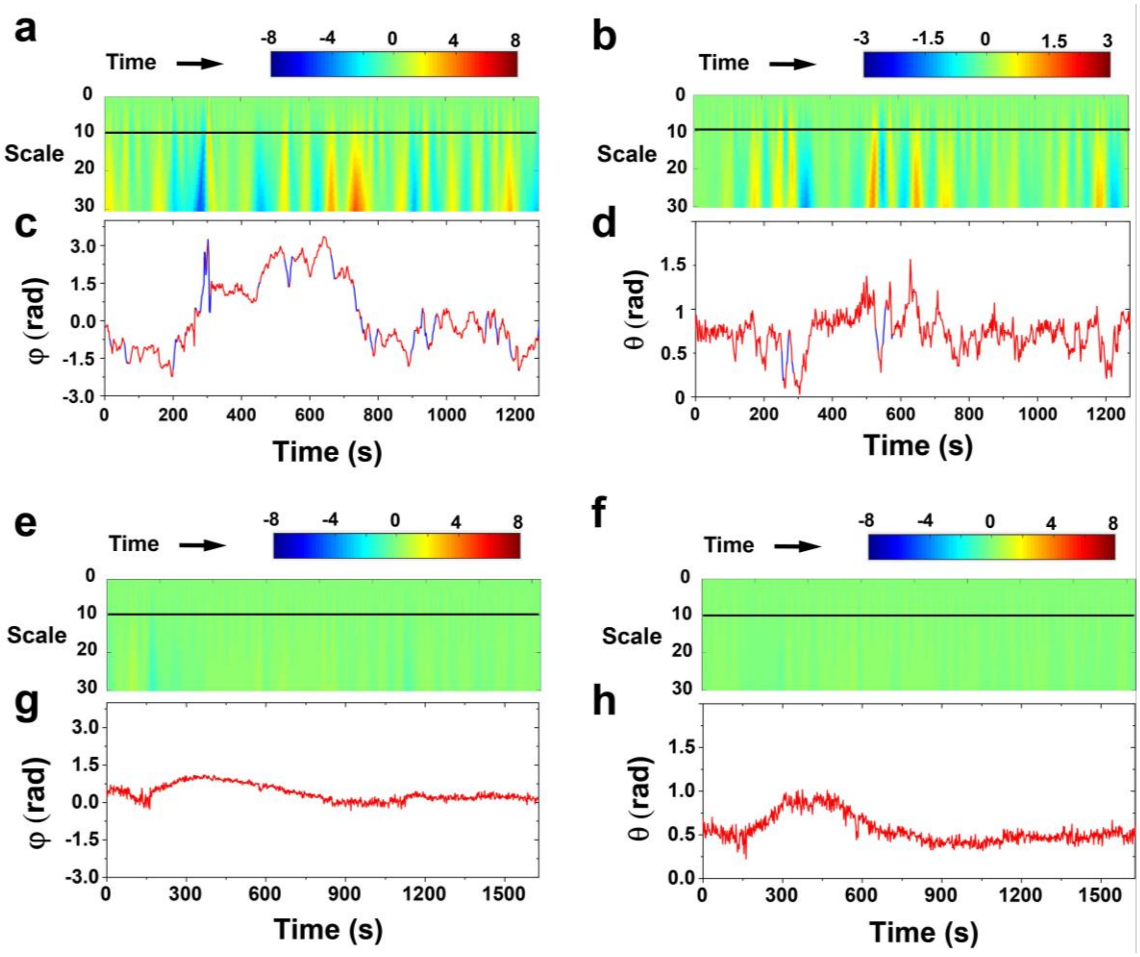


**Supplementary Fig. 8.** Wavelet analysis for distinguishing active rotation from random fluctuations. Color-coded wavelet coefficients obtained from the Haar continuous time wavelet transform of the azimuthal (**a**) and polar (**b**) rotational angle for a single endosome trajectory are plotted against time (x-axis) for different widths of the wavelet function (y-axis), which has the unit of the number of image frames and is referred to as "scale". The black line indicates scale = 10, which was chosen to separate passive and active rotations. The azimuthal (**c**) and polar (**d**) rotational angle of the same particle analyzed in (**a**) and (**b**) are plotted as a function of time. Segments of random noise are shown in red and those of active rotation are in blue. The same wavelet transform analysis was done on the azimuthal (**e**) and polar (**f**) rotational angles of a representative phagosome in a nocodazole-treated microtubule-disrupted cell, and the two rotational angles of this phagosome are plotted in (**g**) and (**h**) as a function of time. No active rotation was detected in the trajectory of this phagosome in a microtubule-disrupted cell. The results are representative of N = 8 phagosomes from 8 microtubule-disrupted cells.

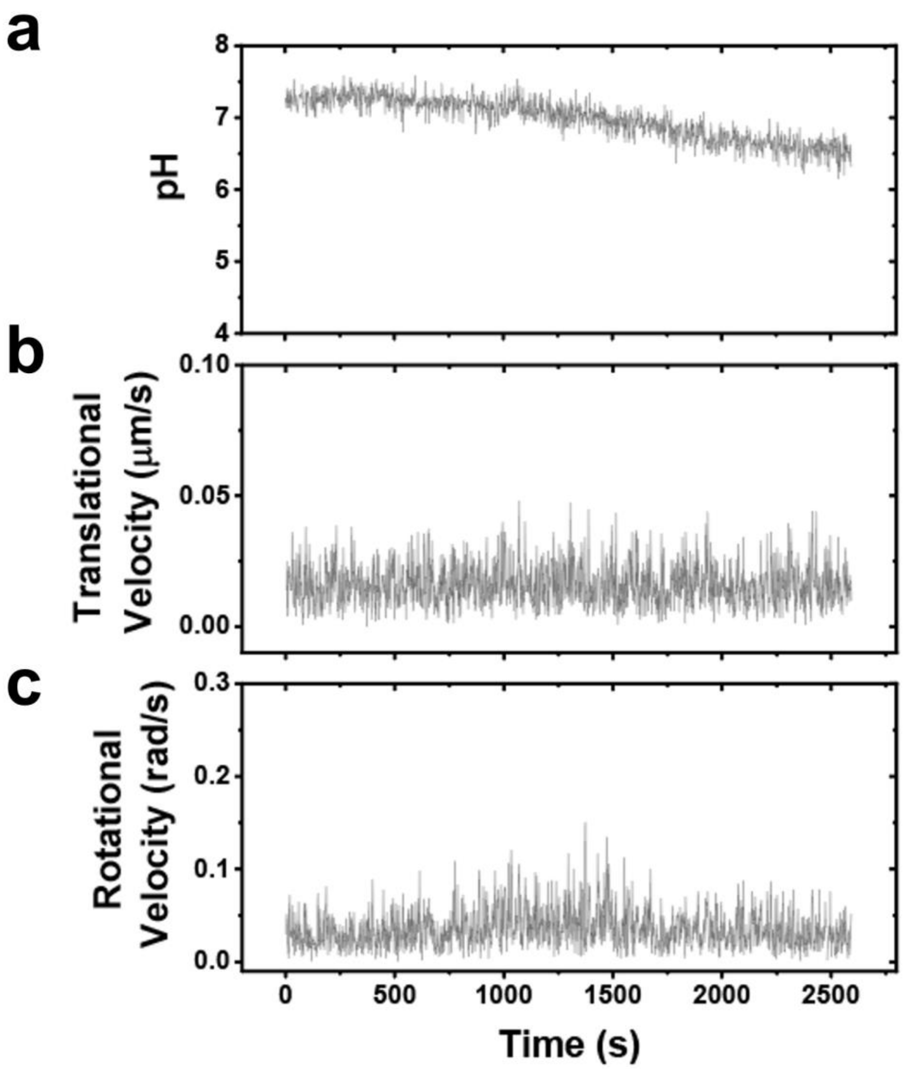


**Supplementary Fig. 9.** (**a-c**) Acidification (**a**), translational velocity (**b**) and rotational velocity (**c**) of a phagosome that never underwent rapid microtubule-based transport. Data are representative of N = 13 phagosomes from 12 cells.

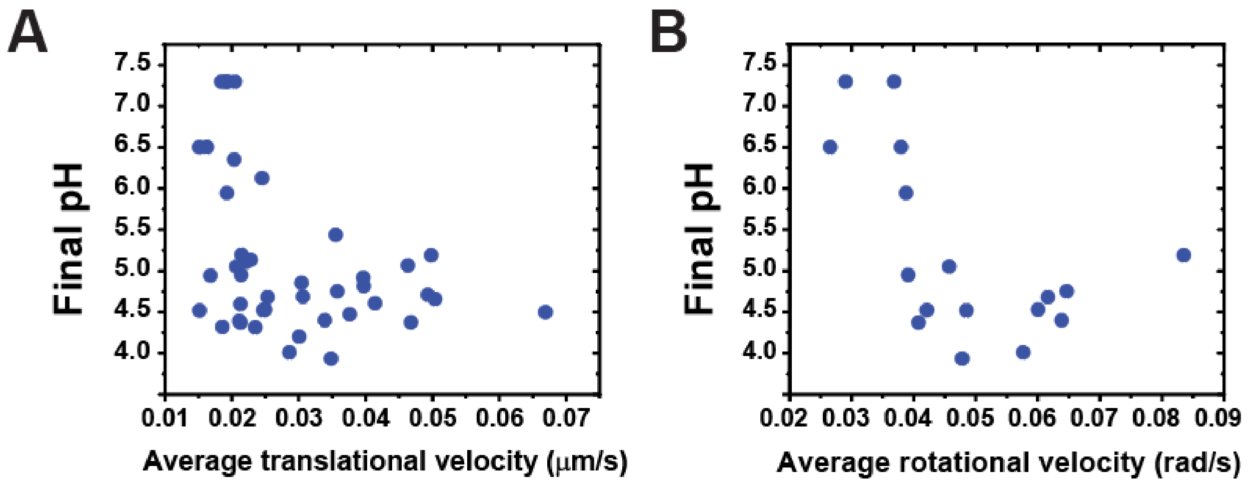


**Supplementary Fig. 10.** (**a-b**) Scatter plots showing acidification rate and final pH of single phagosomes plotted separately against their translational and rotational velocity during rapid acidification period in activated RAW264.7 macrophage cells. For translational tracking, N = 42 phagosomes from 24 cells. For rotational tracking, N = 17 phagosomes from 12 cells.

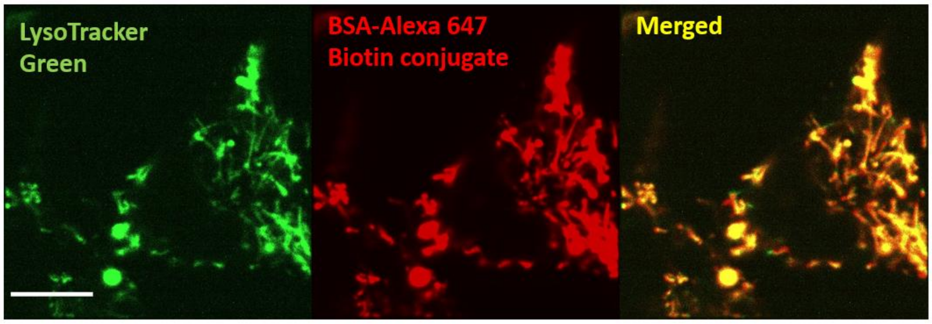


**Supplementary Fig. 11.** Re-scan confocal microscopy (RCM) images showing the colocalization of Lysotracker Green with intracellular organelles containing BSA-Alexa647 in RAW 264.7 macrophage cells. Scale bar, 10 μm.

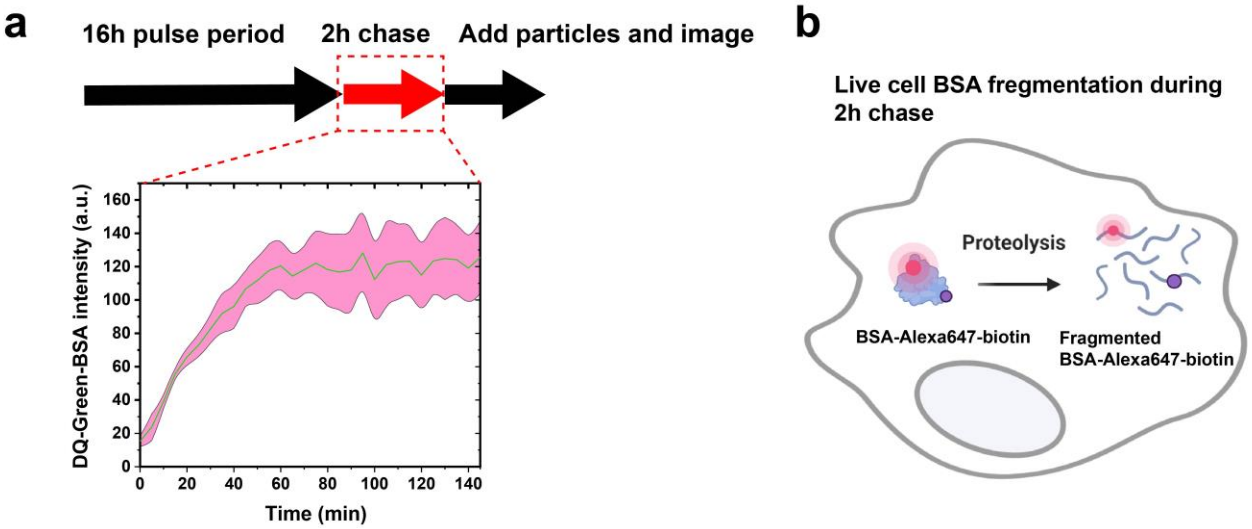


**Supplementary Fig. 12.** Schematics indicating the workflow of the pulse-chase procedure. (**a**) Quantitative analysis of the intensity of endocytosed DQ-green BSA during chasing. Shaded area represents standard deviation from 5 cells (**b**) Schematic illustration of internalized BSA-Alexa647-biotin being proteolyzed and fragmented inside lysosomes during the 2h chase period, which was before FRET-RotSensors were added to cells for internalization.

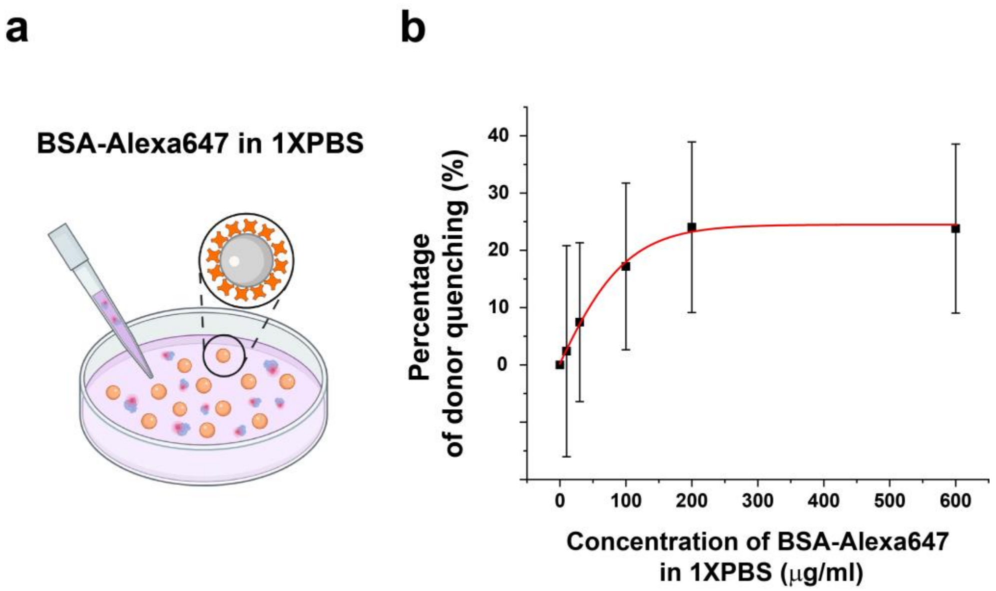


**Supplementary Fig. 13.** *In vitro* control showing FRET between particle-coated donor fluorophore (SAv-Alexa568) and BSA-Alexa647 (acceptor) added in solution. (**a**) Schematic illustration of the experiment workflow. Donor emission (Ex/Em: 561/586 nm) of the FRET-RotSensors was measured in the presence of fluid-phase BSA-Alexa647 in 1×PBS. (**b**) Percentage of the donor quenching was plotted as a function of BSA-Alexa647 concentration in 1×PBS solution. Data is fitted with sigmoidal-Boltzmann function (shown as the red solid line). Error bars represent standard deviation from N = 53 FRET-RotSensors (0 µg/ml), 137 FRET-RotSensors (10 µg/ml), 93 FRET-RotSensors (30 µg/ml), 106 RotSensors (100 µg/ml), 36 RotSensors (200 µg/ml), and 60 FRET-RotSensors RotSensors (600 µg/ml).

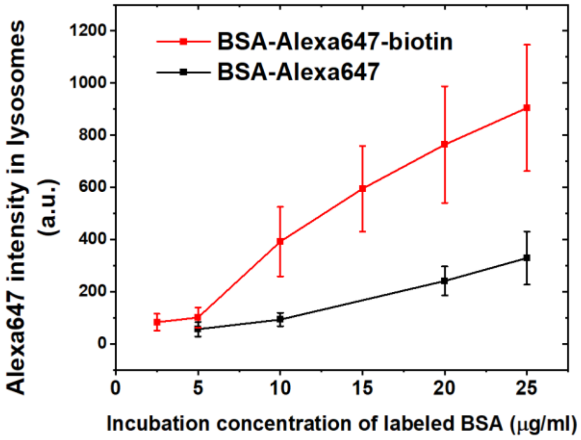


**Supplementary Fig. 14.** Comparison of loading efficiency into lysosomes of BSA-Alexa647-biotin and BSA-Alexa647. Error bars represent standard deviation. BSA-Alexa647 loading data was collected from N = 21 cells (5 μg/ml), 29 cells (10 μg/ml), 19 cells (20 μg/ml) and 34 cells (25 μg/ml). BSA-Alexa647-biotin loading data was collected from N = 16 cells (2.5 μg/ml), 20 cells (5 μg/ml), 22 cells (10 μg/ml), 27 cells (15 μg/ml), 13 cells (20 μg/ml), and 13 cells (25 μg/ml).

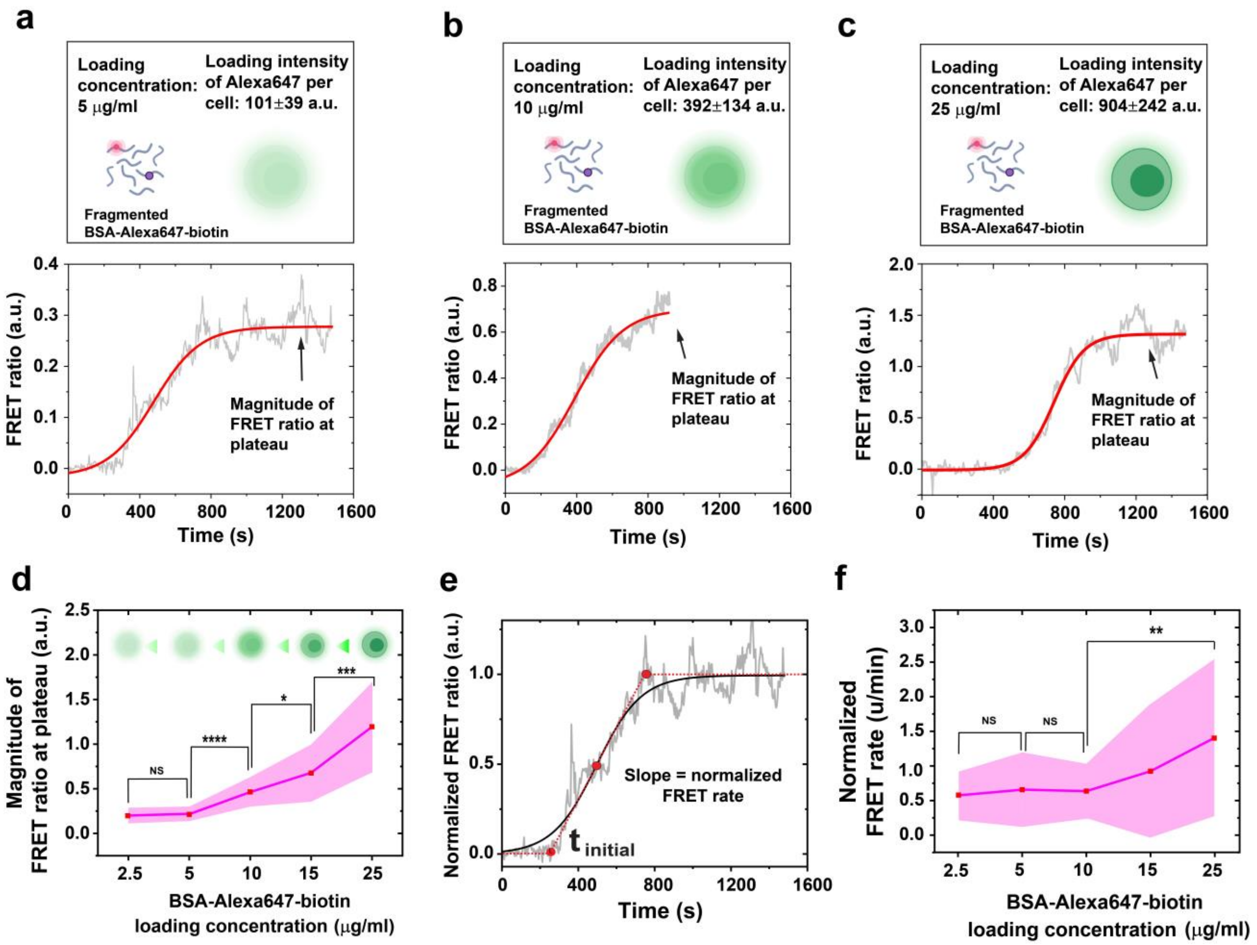


**Supplementary Fig. 15.** Optimization of FRET fusion assay by adjusting loading concentration of BSA-Alexa647-biotin. (**a-c**) Representative FRET profiles obtained from FRET-RotSensors in macrophage cells at different loading concentrations of BSA-Alexa647-biotin. (**d**) Magnitude of FRET ratio at plateau is plotted as a function of loading concentration of BSA-Alexa647-biotin. (**e**) Demonstration of a normalized FRET ratio vs. time plot and the sigmoidal Boltzmann fitting. The slope at the half-response point $t_0$ is defined as normalized FRET rate. (**f**) Normalized FRET rate from individual phagosomes is plotted as a function of loading concentration BSA-Alexa647-biotin. In (**d**) and (**f**), data was collected from N = 18 phagosomes from 14 cells (2.5 μg/ml), 16 phagosomes from 12 cells (5 μg/ml), 19 phagosomes from 12 cells (10 μg/ml), 19 phagosomes from 14 cells (15 μg/ml), and 18 phagosomes from 14 cells (25 μg/ml). Statistical significance is highlighted by p-values (student's t-test) as follows: * $p < 0.05$, ** $p < 0.01$, *** $p < 0.001$, **** $p < 0.0001$, NS $p > 0.05$.

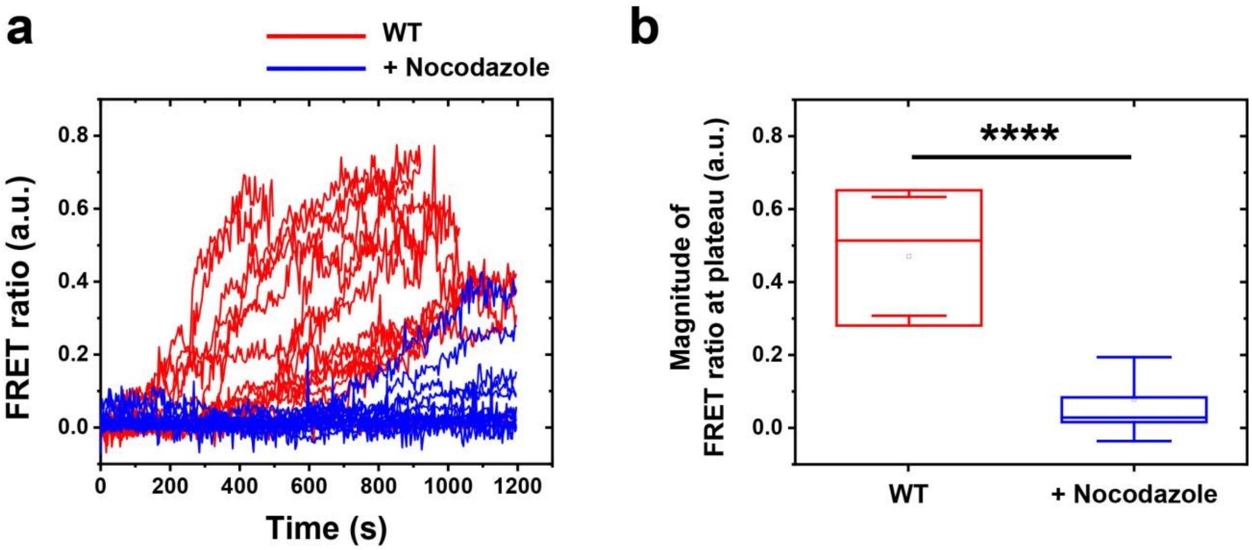


**Supplementary Fig. 16.** Effect of microtubule disruption by nocodazole on phagosome-lysosome fusion. (**a**) FRET ratio of single phagosomes is plotted as a function of time in control cells (WT) and cells treated with nocodazole. Loading concentration of BSA-Alexa647-biotin was 10 μg/ml. (**b**) Comparison of magnitude of FRET ratio at plateau for control cells (WT) and cells treated with nocodazole. Each box plot indicates the mean (horizontal line) and the interquartile range from 25% to 75% of the corresponding data set. Data were collected from N = 19 phagosomes in 12 WT non-treated cells, and 13 phagosomes from 10 nocodazole-treated cells. Statistical significance is highlighted by the p-value (student's t-test) as follows: **** $p < 0.0001$.

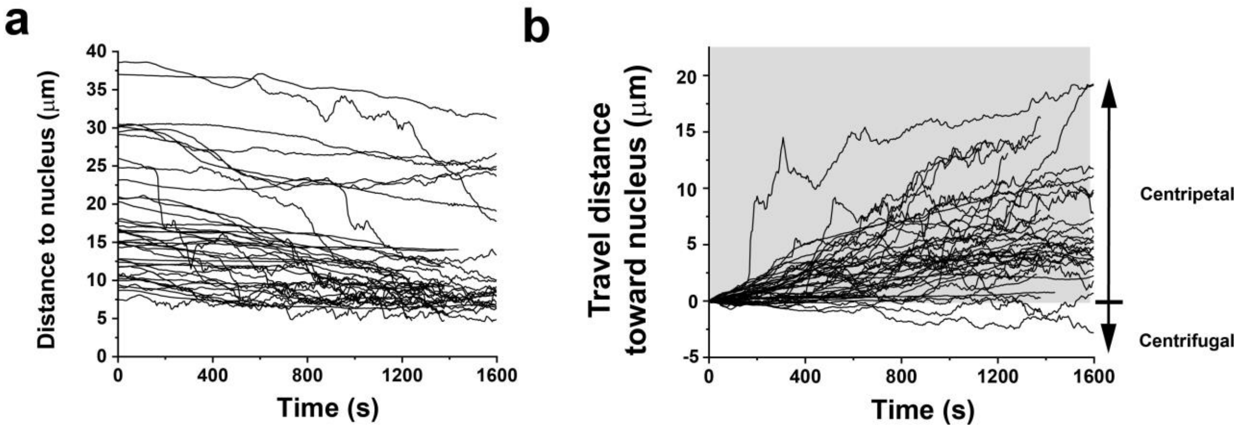


**Supplementary Fig. 17.** Quantification of phagosome centripetal movement. (**a**) The distance of single phagosomes to the centroid of the nucleus is plotted as a function of time. N = 40 phagosomes from 22 cells. (**b**) The displacement of phagosome in (**a**) towards the nucleus center was plotted against time to separate centripetal from centrifugal movement.

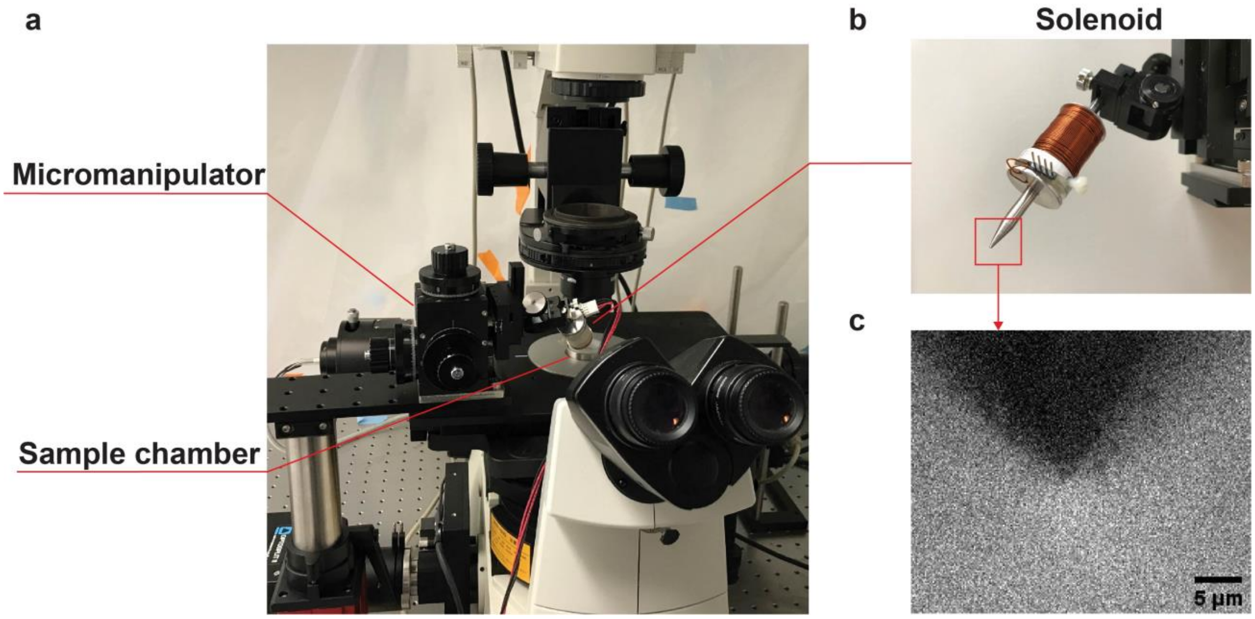


**Supplementary Fig. 18.** Magnetic tweezers setup. (**a**) Picture of the magnetic tweezers setup installed on an inverted Nikon microscope. (**b**) Picture of the solenoid. The metal core is inserted into an aluminum bobbin wrapped with copper coils. (**c**) Bright-field image of the solenoid tip.

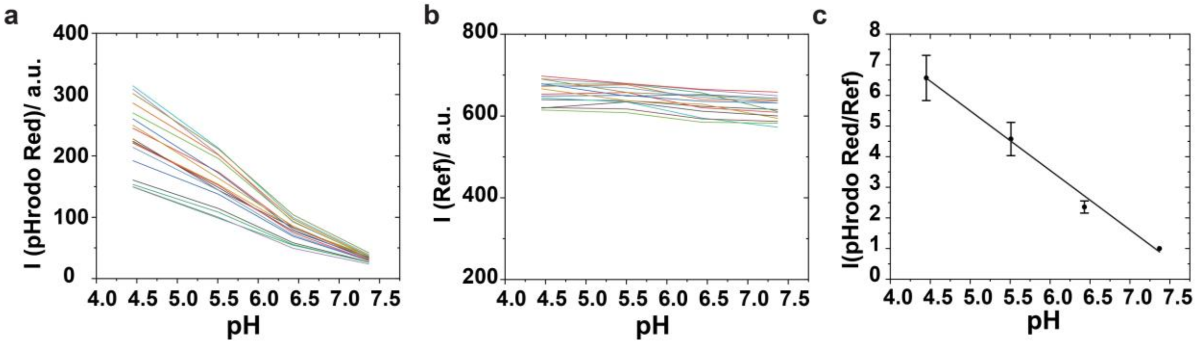


**Supplementary Fig. 19.** Extracellular pH calibration of MagSensors. Fluorescence intensity of pH-indicator pHrodo Red (**a**) and reference dye CF640R (**b**) on individual MagSensors is plotted against pH in aqueous buffer solutions. Each curve represents data from a single MagSensor and a total of 15 particles were analyzed for each sample. (**c**) The fluorescence intensity ratio of pHrodo Red and CF640R of individual MagSensors was calculated and averaged to obtain a pH calibration plot in aqueous buffer solutions. Error bars are standard deviations from 15 individual MagSensors. $R^2$ value of the linear fitting is 0.99.

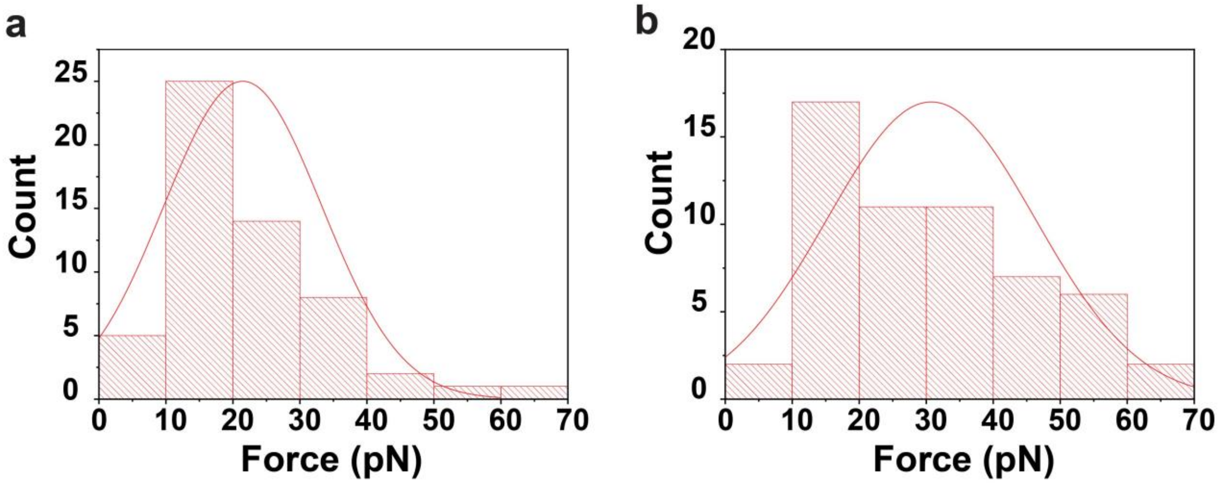


**Supplementary Fig. 20.** Distribution of estimated magnetic force exerted on individual phagosomes at the beginning (**a**) and the end of experiments (**b**). The average magnetic force exerted at the beginning and the end of magnetic pulling was around 21 pN and 31 pN, respectively.

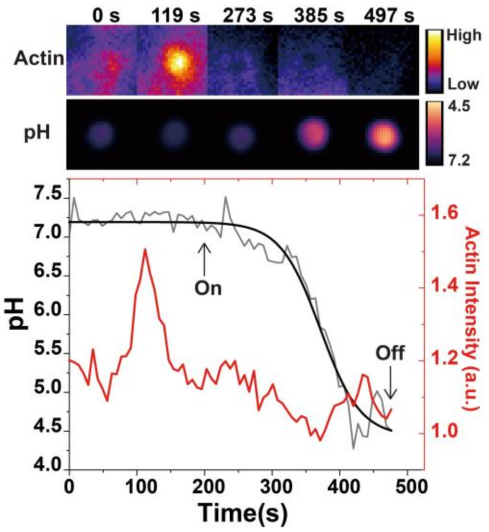


**Supplementary Fig. 21.** Changes of pH and actin intensity of a representative phagosome during acidification under magnetic pulling. Fluorescence images showing the phagosome pH and actin-GFP in cells. pH and intensity of actin-GFP are color-coded based on the scales indicated. Scale bars, 1 µm. The change of phagosome pH and the corresponding sigmoidal-Boltzmann fitting are plotted in black lines. Actin-GFP intensity around the phagosomes is plotted against time in red line. “On” and “off” indicate the start and end of magnetic pulling.

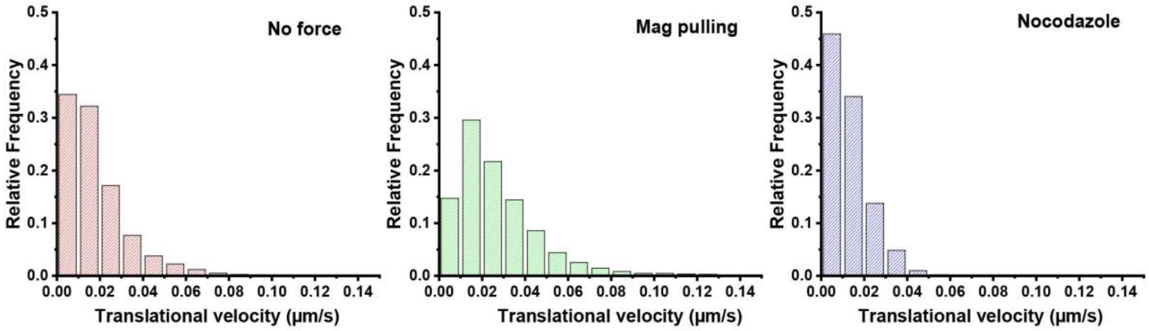


**Supplementary Fig. 22.** Histograms showing the distribution of translational velocities of phagosomes in resting RAW264.7 macrophage cells under different experimental conditions: no force, with magnetic pulling, or after nocodazole treatment. N = 16 phagosomes from 15 cells (no force), N = 17 phagosomes from 16 cells (magnetic pulling) and N = 15 phagosomes from 10 cells (nocodazole).

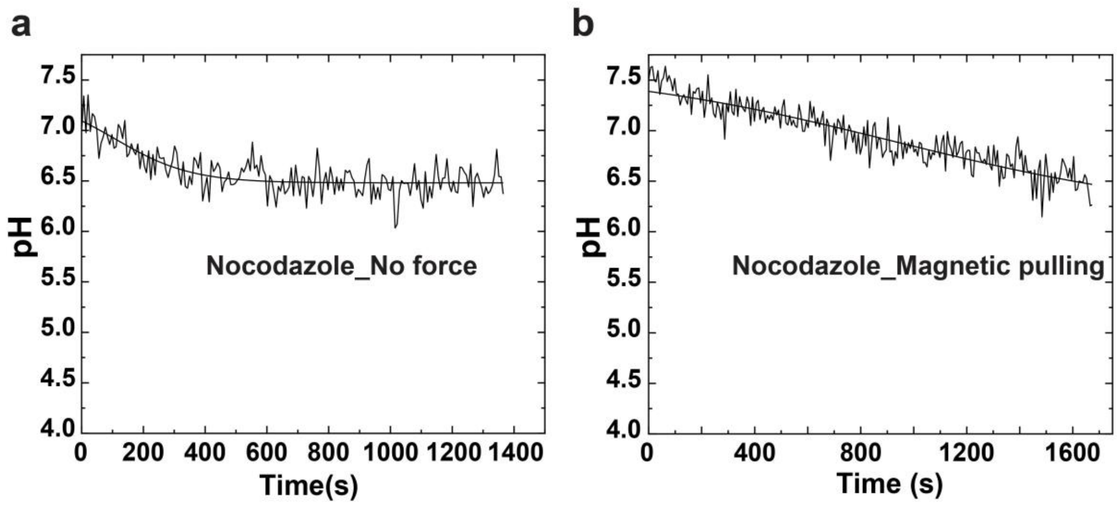


**Supplementary Fig. 23.** Acidification profile of representative single phagosomes after nocodazole treatment without (**a**) or with (**b**) magnetic pulling. Each pH-time curve is fitted with sigmoidal-Boltzmann function. Data are representative of N = 9 phagosomes from 8 cells in (after nocodazole treatment, no force) and N = 10 phagosomes from 7 cells in (after nocodazole treatment, magnetic pulling).

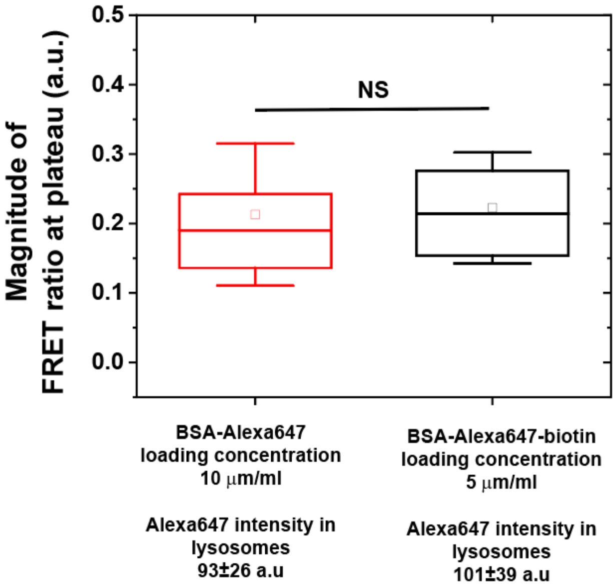


**Supplementary Fig. 24.** Comparison of the magnitude of FRET ratio at plateau (maximum FRET ratio) when BSA-Alexa647 was used instead of BSA-Alexa647-biotin. Different loading concentrations of BSA proteins were used to achieve similar concentration of acceptors in lysosomes (Alexa647 intensity in lysosomes). Data was collected from 14 phagosomes from 10 cells (BSA-Alexa647) and N = 16 phagosomes from 12 cells (BSA-Alexa647-biotin). Statistical significance is highlighted by the p-value (student's t-test) as follows: NS $p > 0.05$.

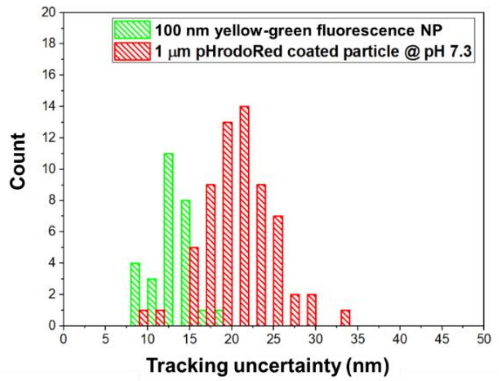


**Supplementary Fig. 25.** Histograms showing localization uncertainty for the 100 nm yellow-green nanoparticle and the 1 μm pHrodoRed-coated particle in each RotSensor.

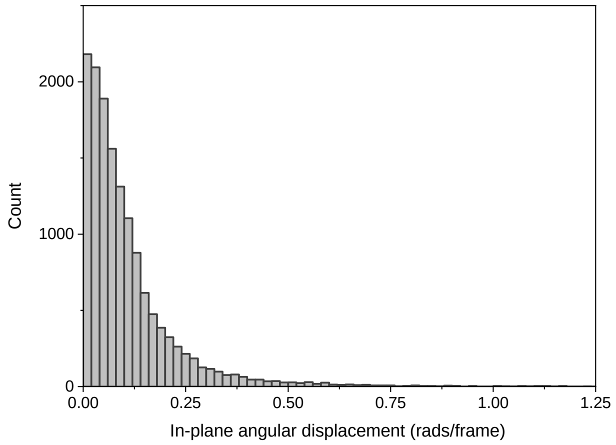


**Supplementary Fig. 26.** Histogram showing the distribution of azimuthal angular displacement (Δφ) between consecutive frames (2 s/frame). Average Δφ is 0.12 ± 0.16 rad (6.9° ± 9.2°). N = 17 phagosomes from 12 cells.

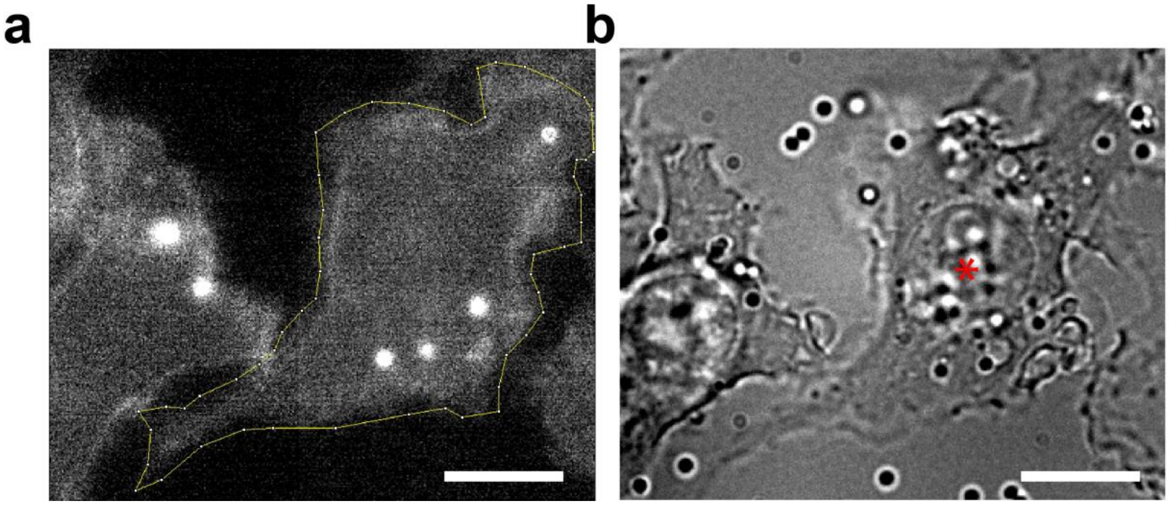


**Supplementary Fig. 27.** Determination of cell boundary and the centroid of cell nucleus center for the centripetal movement analysis. (**a**) Cell periphery is traced based on the autofluorescence emission (ex/em 561/586 nm) of the cell. (**b**) The centroid of the cell nucleus is determined based on bright field images of the cell. Scale bar, 10 μm.

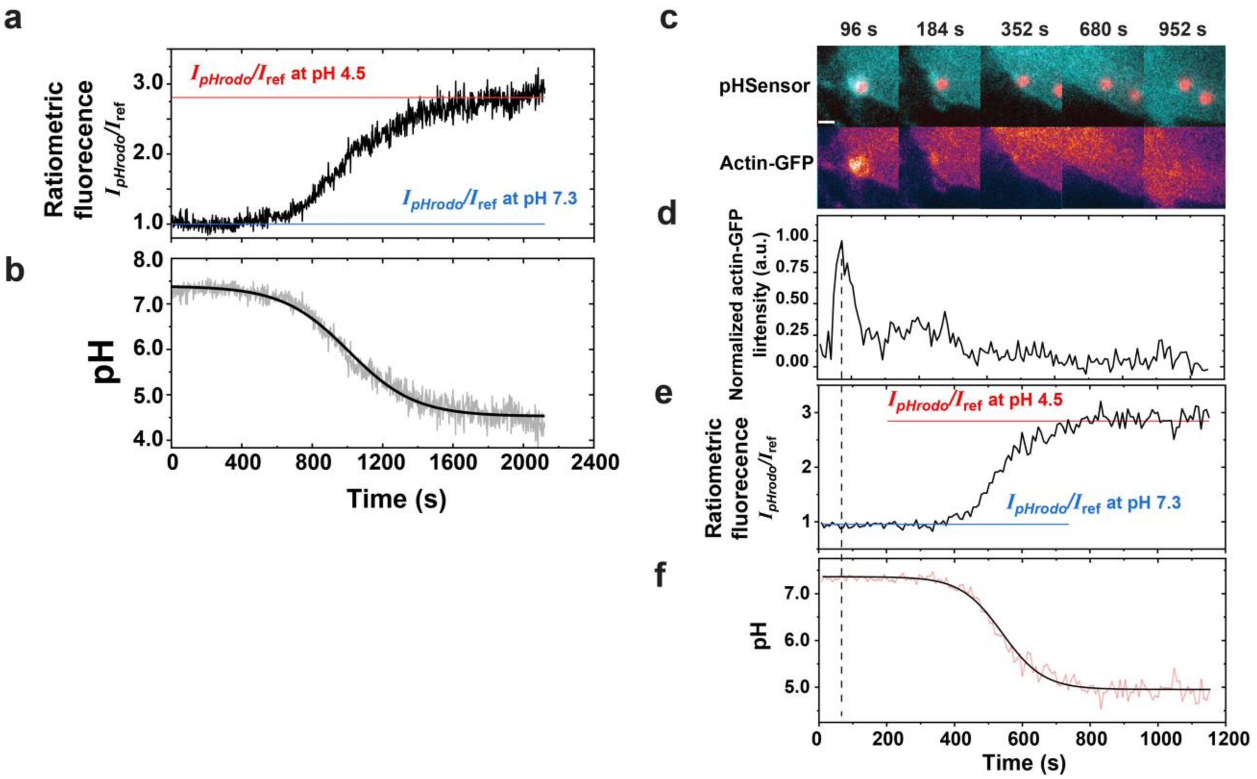


**Supplementary Fig. 28.** (**a**) Ratiometric fluorescence of a representative pH-RotSensor in RAW264.7 phagosome. Horizontal lines indicate ratiometric emission of the pH-RotSensor measured at pH 4.5 and pH 7.3. The ratiometric fluorescence of the pH-RotSensor was converted to pH (**b**). (**c-f**) Phagosome acidification in actin-GFP expressing RAW267.4 macrophage cells. (**c**) Fluorescent images showing subcellular locations of phagosomes containing pH-RotSensors at different time points in actin-GFP expressing cells. Scale bar, 2 µm. (**d**) Normalized actin-GFP intensity around the phagosome (shown in **c**) is plotted as a function of time. (**e**) Ratiometric fluorescence of the pH-RotSensor in (**c**) plotted against time. Horizontal lines indicate ratiometric emission of the pH-RotSensor measured at pH 4.5 and pH 7.3. The ratiometric fluorescence of the pH-RotSensor was converted to pH (**f**).
